# A novel architecture of PT neuron-based corticothalamic connectivity in the auditory system

**DOI:** 10.1101/2023.08.08.552413

**Authors:** Fenghua Xie, Yixiao Gao, Tao Wang, Mengting Liu, Kexin Yuan

**Author notes:** Correspondence (K.Y.), (F.X.). These authors contributed equally (Fenghua Xie, Yixiao Gao).

## Abstract

Largely topographical projections from different modules of the thalamus, such as the primary, secondary and association sensory thalamus, to hierarchically defined cortical areas have been recognized across sensory systems. However, how corticothalamic projections, which are believed to be crucial for the remarkable flexibility and precision exhibited by our sensory systems, are organized remained poorly understood compared with the thalamocortical counterpart. Here we report that, first, the primary auditory thalamus received direct inputs from cortical L5 neurons. Second, in contrast to the robust thalamocortical topography, L5 neurons in each of the primary, secondary and association auditory cortical regions project to each individual module of the auditory thalamus at the macroscale. Third, the association cortex provided the most L5 inputs to all thalamic modules followed by the secondary and primary auditory cortices. Lastly, L5 axon terminals were mainly varicosity-type and evenly distributed across thalamic modules, but those in the polymodal association module were the largest. Our data suggest that all the modules of the auditory thalamus may be under the modulation of common L5 inputs. This fully-connected-like corticothalamic architecture urges a revision of the traditional hierarchical model in the sensory systems.

## Introduction

The bidirectional connection between the sensory thalamus and the sensory cortex plays a crucial role in the functional processing of sensory information in mammalian brains^1–5^. The sensory thalamus is composed of three modules: primary, non-primary and association^6–9^. The primary and non-primary modules receive unimodal and polymodal sensory inputs, respectively, whereas the association module can integrate sensory inputs of multiple modalities with information related to animal’s behavioral states^7,10^. These three modules of sensory thalamus project to the cortex largely in a topographical manner, that is, they preferentially project to the middle layers of the primary, secondary and association sensory cortices^7,11–13^, respectively, although the non-primary and association thalamic neurons may also have axons fibers that extend horizontally in the superficial layer to other cortical areas^7,9,13–15^. These topographically organized thalamocortical pathways serve as essential conduits for sensory integration, perception, and the construction of our conscious experience of the world around us.

The other pathway in the sensory systems that is parallel with the thalamocortical projections is the descending pathway originating from the cortex. Interestingly, although the corticothalamic system has been much less well-understood compared with the thalamocortical system, descending presynaptic terminals greatly outnumber their ascending counterparts at least in the visual and somatosensory thalamus^11,16–18^, suggesting their important contribution to thalamic function. Indeed, it has been shown that the corticothalamic pathways can dynamically modulate thalamic activity^19,20–25^, enabling active and adaptive sensory processing in the thalamus^20,26–28^. However, despite the fact that both layer 5 (L5) and layer 6 (L6) neurons project to the thalamus, most of our knowledge about the organization and function of the corticothalamic descending pathways have actually been obtained from layer 6 projections in the lemniscal sensory systems^3,20,26,29,30^, which provide feedback inputs to the primary sensory thalamic nuclei.

Pyramidal neurons in L5 are among the most extensively studied cortical neurons, and their morphology, connectivity patterns and intrinsic electrophysiological properties have been closely examined across species^31^. Based on the difference in projection pattern, L5 neurons can be classified into two types, the intratelencephalic (IT) type and the pyramidal tract (PT) type. L5 IT neurons are characterized by their projections almost restricted within the cortex, whereas L5 PT neurons project to a number of subcortical structures including the thalamus. It is believed that L5 PT neurons in lower-order sensory cortices provide feedforward-type inputs selectively to higher-order sensory thalamus, which in turn relay the received information to higher-order sensory cortices^3,13,32–34^, known as the transthalamic circuit model^35^. Nonetheless, in the sensory systems, the organization and function of L5 projections to the thalamus have received much less attention compared with those to other subcortical structures such as the striatum^36–38^ and midbrain^39–43^, probably due to the sparse labeling of L5 neurons when retrograde tracers, including biotinylated dextran amin (BDA)^13^, cholera toxin subunit B (CTB)^44^ and glycoprotein-deleted rabies viruses (RV)^7,45^, were injected into the sensory thalamus. Intriguingly, some recent studies showed that rAAV2-retro, a viral tool for retrograde tracing, can efficiently label L5 neurons^46^. If rAAV2-retro was used to characterize the corticothalamic (CTh) pathways, to what extent the sparse labeling of L5 neurons and the lower-order L5→higher-order thalamus innervation preference would still hold remains unclear.

To address the above question, we performed 3D modeling of the auditory thalamus for precise anatomical data registration, injected rAAV2-retro into the auditory thalamus of wild-type or transgenic mice to retrogradely label CTh neurons, used double-recombinase system-enabled circuit-specific sparse labeling strategy to depict corticothalamic circuits, applied our newly developed rapid and deformation-free on-slide tissue-clearing method to facilitate high-quality imaging, conducted high-throughput and calibrated measurement of terminal size to compare the morphological property of axon terminals, and performed immunofluorescent staining to determine CTh cell types. We found that, in contrast to the lower-order L5→higher-order thalamus proposal, each individual subdivision of the AT, including the primary auditory thalamus, receives direct inputs from a large number of L5 PT neurons, which are, notably, distributed in both the primary and higher-order auditory cortices and temporal association cortex (TEa). Interestingly, the TEa provides the most thalamus-projecting L5 PT neurons for all AT subdivisions. Furthermore, the density of L5 axonal terminals, which are predominantly varicosity-type, is similar across AT subdivisions, but the size is significantly larger and more heterogeneous in the polymodal association AT, which was further verified by selectively labeling L5 PT neurons using mscRE4 enhancer virus^47^. Systematic comparisons demonstrated considerable distinctions between the thalamic projection patterns of L5 PT and L6 CT neurons.

## Results

### The primary auditory thalamus receives strong inputs from cortical layer 5

To accurately register anatomical data to subdivisions of the auditory thalamus (AT), we first conducted a 3D modeling of the AT followed by the construction of a 2D coronal atlas, which was based on YFP and CB immunostaining signals in Thy1-YFP-H mice ^48–51^ (Fig S1 and Methods). Next, we took advantage of viral tracing tools ^52^ to retrogradely label cortical neurons projecting to the AT. We locally injected rAAV2- retro-Cre-eGFP into the MGBv of wild-type mice to express Cre in upstream neurons, then injected AAV9-FLEX-tdTomato into the AUD to label MGBv-projecting AUD neurons (Fig 1A). We observed tdTomato-labeled both L5 and L6 neurons in the AUD when the expression of eGFP was localized in the MGBv (Fig 1B, Fig S2A). The depth and laminar distribution of these neurons were verified by registering them to the Allen Mouse Atlas^44,53,54^ (Fig 1C, FigS2B). Labeled L5 neurons were much more numerous and widely distributed in the AUD than labeled L6 neurons (Fig 1D). Since it has been known that the secondary sensory thalamic nuclei^6,55–57^, including the secondary auditory thalamus^58^, receive direct inputs from cortical L5 neurons, our data indicated that L5 neurons project to each individual subdivision of the auditory thalamus but not only to the secondary subdivisions.

**Figure 1.**
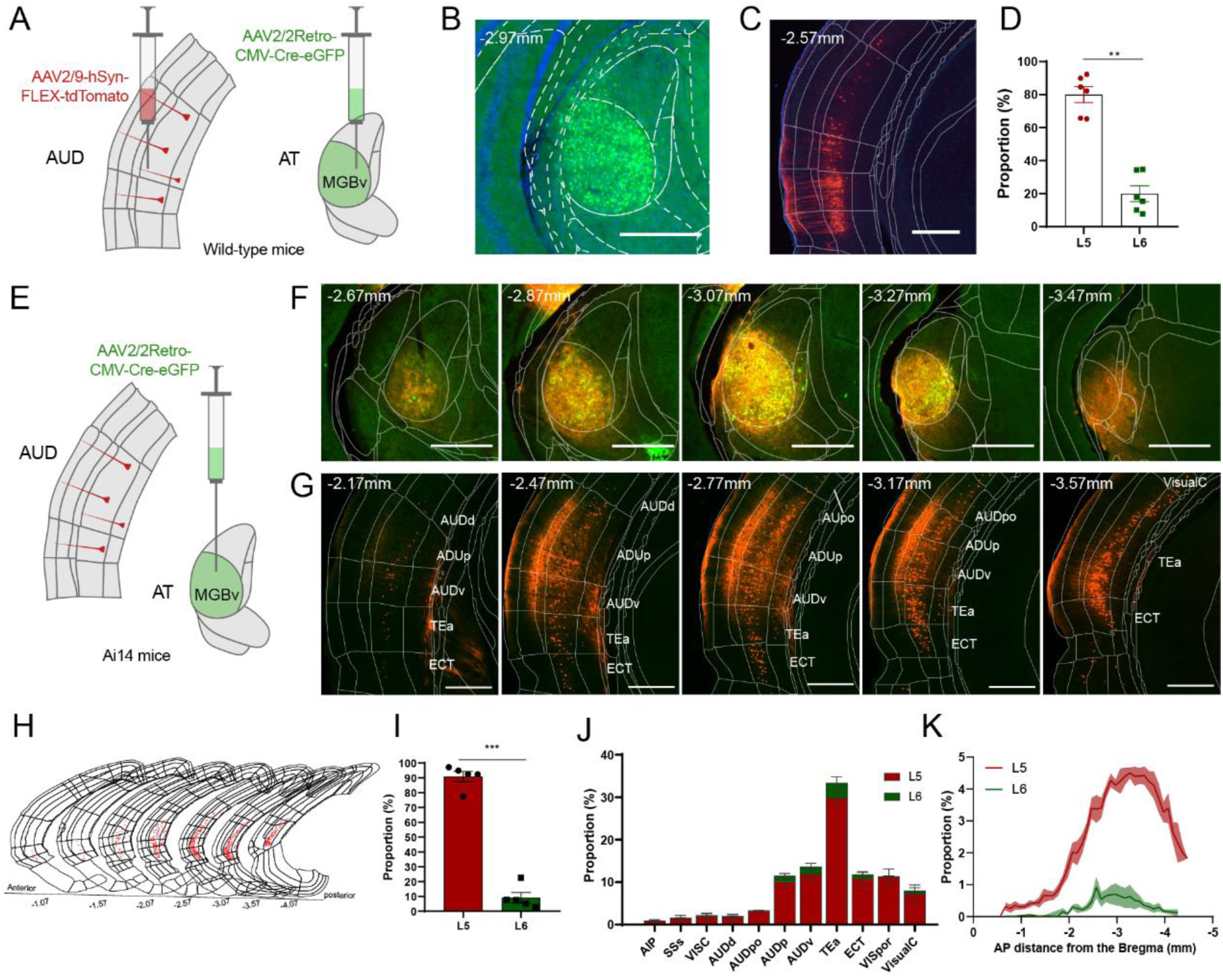
The primary auditory thalamus receives strong inputs from cortical layer5. (A) Schematic diagram of the dual-virus injection experiment for retrograde tracing of MGBv in wild-type mice. (B-C) Representative images of MGBv (B) locally transfected by rAAV2-retro and the auditory cortical areas expressing tdTomato(C) obtained from coronal brain slices. Scale bars, 500μm. (D) Proportion of retrogradely labeled MGBv-projecting neurons in different cortical layers of mice auditory cortex. Data were from 6 mice. L5, 80.0% ± 4.39%. L6 20.0%±4.39%. (E) Schematic representation of the rAAV2-retro injection into the MGBv of transgenic Ai14 mice. (F-G) Representative coronal brain slices showing MGBv transfected by rAAV2-retro (F) and neurons in auditory cortical areas expressing tdTomato (G). Scale bars, 500μm. (H) Overall distribution of MGBv-projecting neurons in the auditory-related cortical areas retrogradely labeled after rAAV2-retro injection into MGBv. (I) Laminar distribution of retrogradely labeled MGBv-projecting neurons in the whole auditory-related cortices. Data were from 5 mice. L5: 90.9%±3.89%; L6: 9.1%±3.89%. (J) Statistical comparison between the proportions of L5 and L6 MGBv-projecting neurons across different cortical regions retrogradely labeled by rAAV2-retro. (K) Anteroposterior (AP) distribution of retrogradely labeled L5 MGBv-projecting and L6 MGBv-projecting neurons in Ai14 mice. Green line represents L6, while red line represents L5. Statistic methods used both in D and I were paired t test. *P* value: \*\**P*<0.01, \*\*\**P*<0.001. More details in TableS2.

Considering that L5 neurons in other cortical regions may also project to the MGBv, we next characterized the distribution of MGBv-projecting L5 (L5→MGBv) neurons in the entire cortex. To do so, we injected rAAV2-retro-Cre-eGFP locally into MGBv of Ai14 mice (Fig 1E, F), which would express tdTomato in all neurons retrogradely infected by rAAV2-retro. We found that L5→MGBv neurons were mainly distributed in all auditory cortical subregions and temporal association cortices, including the AUDd, AUDpo, AUDp, AUDv, TEa, and ECT (Fig 1G-H). Our data showed that rL5→MGBv neurons considerably outnumbered L6 neurons in different cortical regions, at different AP positions, and total numbers (Fig 1I-K). The immunostaining of rL5→MGBv neurons with NeuN showed that they represented about a quarter of L5 neurons (Fig S2E-F). Since a small volume of the virus tended to leak into the hippocampus, which is located right on top of the AT, we intentionally injected the virus into the hippocampus to examine the distribution of hippocampus-projecting cortical neurons. We found that very few neurons in the AUD were labeled, but a substantial number of neurons were observed in the entorhinal cortex (Fig S2G), which makes strong projections to the hippocampus^59^. Thus, the virus leaked into the hippocampus was very unlikely to contribute to retrograde labeling of auditory cortical L5 neurons.

The widely accepted corticothalamic circuit model nowadays is mainly based on data from rats^60^, cats^61,62^, and non-human primates^63^. To determine whether the L5→MGBv circuit is specific to mice, we injected rAAV2-retro-Cre-tdTomato and AAV2/9-FLEX-eGFP into the MGBv and AUD of rats, respectively (Fig S3A-B). Retrogradely labeled neurons were observed both in the L5 and L6 of rat AUD (Fig S3B, right), but the number of L5 neurons was about three times of that of L6 neurons (Fig S3C). These data showed that the L5→MGBv circuit is present in rats as well.

Would cortical L5 neurons in other sensory modalities also project to the corresponding primary sensory thalamus? Toward this end, we locally injected rAAV2- retro-Cre-eGFP into the primary visual (dLGN, Fig S3D, left) or somatosensory (VPM, Fig S3G, left) thalamus of mice. As revealed in the auditory system, a large number of retrogradely labeled L5 neurons were observed in both visual and somatosensory cortices (Fig S3D, right, Fig S3E-G. Fig S3G, right, Fig S3H-I).

Taken together, we provided direct evidence for the presence of L5→primary thalamic nucleus projections originating from a wide range of sensory cortices across sensory modalities and, likely, species.

### The temporal association cortex provides the most layer 5 outputs to each individual module in the auditory thalamus

It has been known that the secondary auditory thalamus receives cortical L5 inputs^58^, how would these L5 neurons be distributed in the cortex compared with rL5→MGBv neurons? To this end, we injected rAAV2-retro-Cre-eGFP into the MGBd or polymodal association nuclei of the auditory thalamus (PoA, including the SG, MGBm, POL, PIN and PP) of Ai14 mice (Fig S4A and Fig S4E). We found that, interestingly, the patterns of both laminar (Fig S4B and F, Fig 2A) and regional distribution (Fig S4D and H, Fig2B-C) of retrogradely labeled L5 neurons were very similar to those of rL5→MGBv neurons. Specifically, the majority of rL5→AT neurons were located in middle L5 (Fig 2A), and they were observed across the entire auditory cortices and temporal association cortex at AP direction (Fig 2B). Regarding regional distribution, about 30% of rL5→AT neurons were distributed in the TEa, and about 10% in the AUDp, AUDv and ECT, respectively (Fig 2C, J). Therefore, the TEa is the cortical region that has the most neurons protecting to each individual subdivisions of AT (Fig 2C).

**Figure 2.**
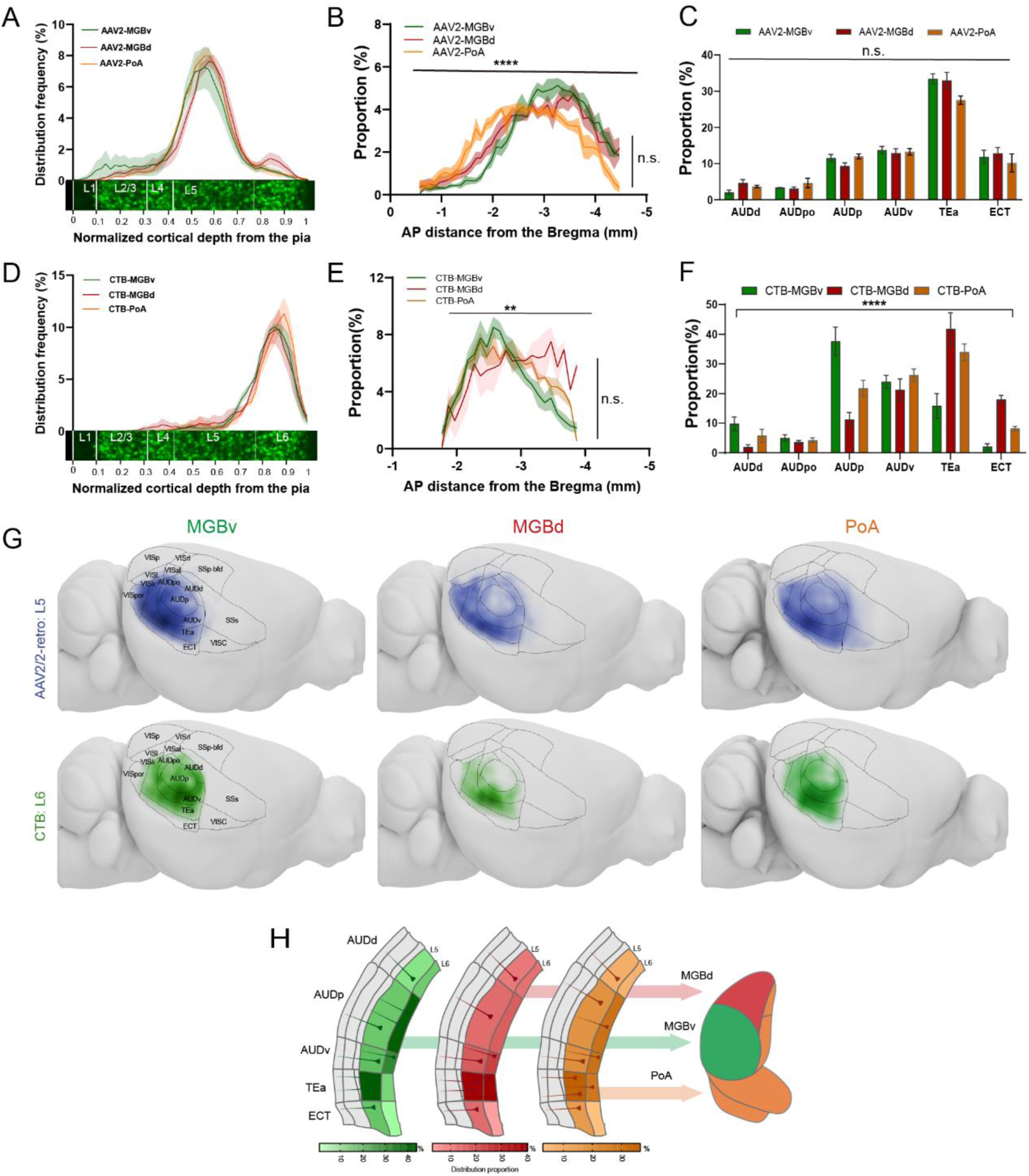
Mapping and distribution comparisons of auditory L5 and L6 neurons projecting to different modules of the auditory thalamus. (A-C) Distribution of neurons retrogradely labeled by rAAV2-retro projecting to the MGBv (green line), MGBd (red line), and PoA (orange line) along the depth from the pia (A, with the pia set at 0 and white matter at 1.), at different AP distances from the Bregma (B), and in different auditory subcortical areas (C). Data were from 6,6 and 7 mice for MGBv, MGBd and PoA, respectively. The shadow areas represent mean±SEM in (A-B). (D-F) Distribution of neurons retrogradely labeled by CTB projecting to the MGBv (green line), MGBd (red line), and PoA (orange line) along the depth from the pia (D), at different AP distances from the Bregma (E), and in different auditory subcortical areas (F). Data were from 5,3 and 3 mice for MGBv, MGBd and PoA, respectively. The shadow areas represent mean±SEM in (A-B). (G) Heatmaps of density distribution estimation for MGBv-projecting neurons (left panels), MGBd-projecting neurons (middle panels), and PoA-projecting neurons (right panels) retrogradely labeled by rAAV2-retro (top panels) and CTB (bottom panels). Dark lines represent borders between adjacent cortical areas. (H) Summarized auditory corticothalamic circuits emphasizing the areal distribution patterns of L5 and L6 neurons projecting to the MGBv, MGBd, and PoA. The projection map in red color represents the quantified density estimation of MGBd-projecting neurons, the green represents MGBv-projecting neurons, and the orange represents PoA-projecting neurons in the auditory cortex, TEa and ECT. L5 and L6 data here were obtained from the averaged proportion of rAAV2-retro and CTB groups across different cortical areas. Statistic methods used in B and E were ordinary Two-way ANOVA, and in C and F were ordinary Two-way ANOVA with Tukey’s multiple comparisons. *P* value, n.s. *P*>0.05, \*\**P*<0.01, \*\*\*\**P*<0.0001. More details in Table S2.

Since RV and CTB have been widely used to map inputs, we also locally injected either of them into the different subdivisions of the AT to examine the potential differences among the three types of tracers in retrogradely labeling corticothalamic neurons (Fig S5A-B and G-H, G&H, M&Q, U&V). Distinct from rAAV2-retro, both RV and CTB preferentially labeled middle L6 neurons in all auditory cortices and the temporal association cortices (RL6→AT neurons, Fig S5C-F, I-L, N-P, R-T, V-X, and Z-AB; Fig 2D), which is consistent with the classic corticothalamic feedback circuits arising from L6^1^ and confirmed that secondary AT receives cortical projections from both L5 and L6^64^. Different from the thalamic subdivision-independent distribution of rL5 neurons, the number of RL6→MGBv and RL6→MGBd neurons was the largest in the AUDp and TEa, respectively, although a similar number of RL6→PoA neurons were observed in the TEa, AUDv, and AUDp (Fig 2E-F). Thus, the distribution pattern of RL6→AT neurons was thalamic subdivision-dependent.

To more accurately characterize the topographic distribution of rL5→AT and RL6→AT neurons, we adopted a Kernel Density Estimation (KDE) method to obtain distribution density estimation of AT-projecting neurons (see Methods for a detailed description of KDE) (Fig S6). In addition to the regional distribution pattern described earlier, we found that rL5→AT and RL6→AT neurons were not evenly distributed in an individual cortical region. Instead, they formed high-density clusters, which largely surrounded the AUDp (Fig 2G).

Collectively, we demonstrated that all the major modules of the auditory thalamus received the most rL5 inputs from the TEa followed by the secondary auditory cortex.

### Layer 5 neurons innervate all auditory thalamic modules with similar axon terminal densities

To infer to what extent corticothalamic inputs may influence the activity in different subdivisions of the AT, we next characterized the distribution of corticothalamic terminals from L5 or L6 in the AT. Toward this end, we injected rAAV2-retro-Flp into the AT to retrogradely express Flp in the cortex, and injected a mixture of diluted rAAV2/9-fDIO-Cre and rAAV2/9-FLEX-Synaptophysin-mRuby-mGFP into temporal cortices to enable Flp-assisted (Fig 3A; see Methods for more details of viral injection strategy), relatively sparse labeling of AT-projecting neurons (Fig 3B-D, E1-E3, F1-F3) and their terminals in the AT (Fig 3E4-E5, F4-F5), which made the identification and quantification of axon terminals more accurate. Furthermore, we developed a rapid and non-scaling on-slide tissue clearing method (RSOTC; Fig S7A-B, see Methods for more details) to facilitate higher-quality optical imaging of anatomical structures including axons and terminals (Fig S7C). This method enabled the confirmation of synaptic connections between auditory AT-projecting neurons expressing Synaptophysin-mRuby^65–67^ and AT neurons expressing PSD95-FingR-eGFP^68,69^ (Fig S7D).

**Figure 3.**
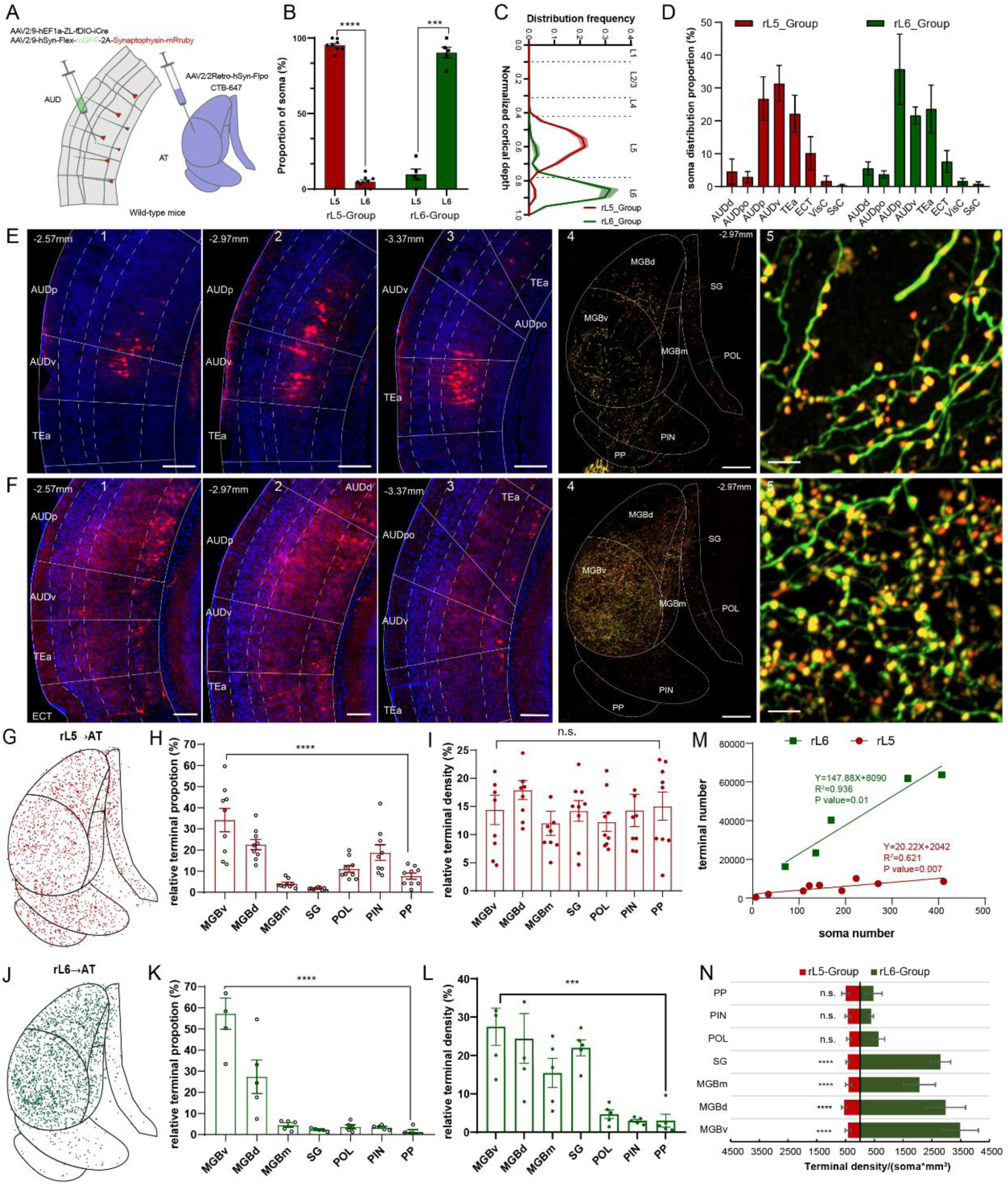
Differential distribution patterns of axon terminals in the AT between retrogradely labeled L5→AT and L6→AT neurons. (A) Schematic diagram illustrating the labeling of auditory corticothalamic neurons and their axon terminals with mGFP-2A-Synaptophysin-mRuby by combining rAAV2-retro injection into the auditory thalamus and the double recombinase Cre and Flip systems. (B) Fractions of L5 and L6 neurons among selectively labeled L5 and L6 AT-projecting neurons (rL5-group and rL6- group), respectively. (C) Depth distributions of L5 and L6 AT-projecting neurons along the normalized cortical depth, with the pia set at 0 and white matter at 1. The red line with the corresponding shadow indicates L5 AT-projecting neurons, while the green line with the corresponding shadow indicates L6 AT-projecting neurons. The dotted line represents the boundaries between adjacent layers, determined by the cytoarchitecture of the auditory cortex. Data for the rL5- group were obtained from 9 hemispherical brains of 5 mice, while data for rL6-group were obtained from 5 hemispherical brains of 5 mice. Data presented next are the same sources for both groups. (D) Proportions of retrogradely labeled L5 and L6 AT-projecting neurons in different auditory-related cortices and adjacent cortical areas, such as the visual or somatosensory cortex. (E-F) Representative confocal images of AT-projecting neurons only distributed in L5 (E) or L6 (F), respectively, of auditory-related cortical areas (1-3), axon terminals of auditory L5 or L6 neurons in the auditory thalamic nuclei including MGBv, MGBd, MGBm, SG, POL, PIN and PP (4) and zoom-in details of axon terminals located in the MGBv (5). Blue represents DAPI. Red in panels E1-E3 and F1-F3 represent L5 AT-projecting neurons and L6 AT- projecting neurons, respectively. Green in panels E4-E5 and F4-F5 represent axons derived from L5 AT-projecting neurons and L6 AT-projecting neurons, respectively. Merged green and red in panel E4-E5 and F4-F5 represents axon terminals from L5 AT-projecting neurons and L6 AT-projecting neurons, respectively. Scale bars in (E1-E3) and (F1-F3), 200μm. Scale bars in E4 and F4, 150μm. Scale bars in E5 and F5, 5μm. (G-L) 3000 weighted points of axon terminals from all L5-group samples (G) or all L6-group samples (J) were mapped in the auditory thalamus. Proportions and density distributions of axon terminals of retrogradely labeled L5 AT-projecting neurons (H-I) and L6 AT-projecting neurons (K-L) in different subnuclei of the AT. Data are presented as mean±SEM. (M) Correlation between retrogradely labeled AT-projecting neurons in the rL5-group and rL6-group and their terminals in the auditory thalamus. Red solid circles and corresponding linear regression line represent rL5-group. Green solid squares and corresponding linear regression line represent rL6-group. The goodness of fit is represented by R^2^ in the corresponding color. The slope significance of non-zero is represented by the P value in the corresponding color. (N) Terminal density per volume for each neuron of the retrogradely labeled rL5-group and rL6-group AT-projecting neurons. RM one-way ANOVA with Dunnett’s multiple comparisons tests were performed in H, K and L, and Friedman test with Dunn’s multiple comparisons test was used in I. For comparing of the same AT subdivision between rL5 and rL6 in N, multiple unpaired t tests with correction for using the Holm-Sidak method was performed. P value, n.s. *P*>0.05, *** *P*<0.001, **** *P*<0.0001. More details in TableS2.

Considering that the size of different auditory thalamic subdivisions varies, the number of corticothalamic terminals in a certain subdivision may not be able to appropriately suggest the strength of cortical influence. Therefore, we calculated the relative density of corticothalamic terminals instead in each thalamic subdivision (see Methods for more details of relative density calculation). Interestingly, the relative density of rL5→AT axon terminals was not significantly different across different subdivisions, including the MGBv, MGBd, MGBm, SG, POL, PIN, and PP, although the number of terminals varies (Fig 3G-I, Fig S8A-D). In contrast, the density of rL6→AT axon terminals was significantly higher in the MGBv, MGBd, MGBm, and SG than that in the POL, PIN, and PP (Fig 3J-L, Fig S8E-H), suggesting that the MGB is under stronger modulation of cortical feedback inputs^70–74^. We further found that the density of axon terminals in the MGBv, MGBd, MGBm and SG from L6 was about seven times higher than that from L5, which could be accounted for by a considerably higher estimated number of terminals contributed by single L6→AT neurons (Fig 3M- N; see Methods for more details of estimation).

Collectively, axon terminals of rL5→AT neurons demonstrated thalamic subdivision- independent density distribution, whereas those of rL6→AT neurons were more numerous and denser in the principal nucleus of the AT.

### Layer 5 neurons send axons with larger terminals to the association module of the auditory thalamus

After characterizing the density distribution of cortical axon terminals in the AT, we conducted a closer examination of these terminals in terms of their morphological features and their distribution along axons (Fig 4A-B; See Methods for more details of imaging and image analysis techniques).

**Figure 4.**
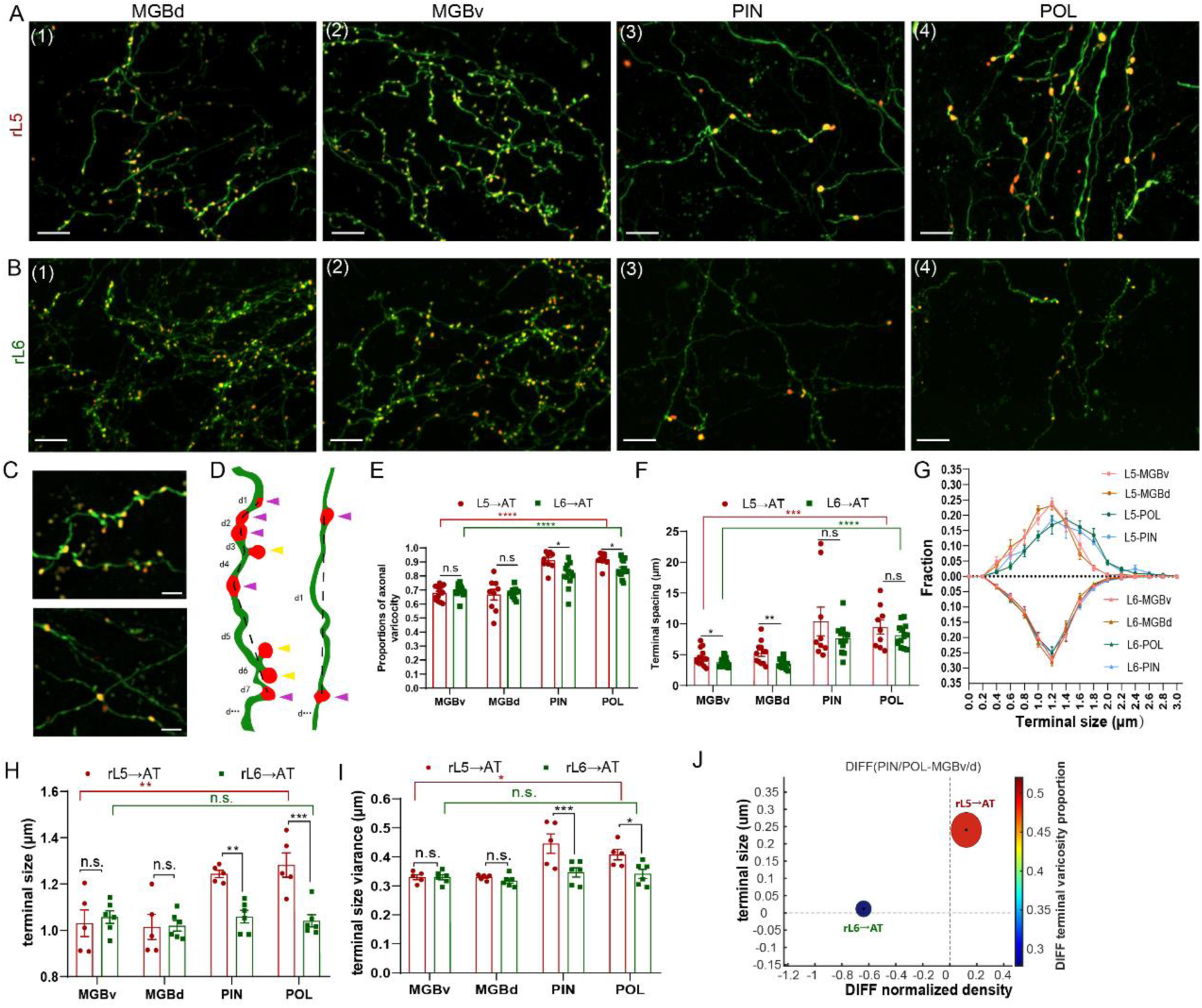
Characteristics of axon terminal morphology in the auditory thalamus derived from auditory L5 and L6 neurons. (A-B) Representative confocal images displaying axons (green) and terminals (overlap of green and red) distributed in MGBv (A1 and B1), MGBd (A2 and B2), PIN (A3 and B3), and POL (A4 and B4) from L5 AT-projecting neurons(A) and L6 AT-projecting neurons in the temporal cortices. Scale bars in (A-B), 10μm. (C) Enlarged views of corticothalamic axon terminals from auditory cortical AT-projecting neurons. Top panel shows two axon fibers with a dense distribution of terminals, while bottom panel shows three axon fibers with sparse terminals. Scale bars, 4μm. (D) Sketches of two axon fibers with different morphologies and terminal distributions showing classification of axon terminals and measurements of axon terminal spacing. Green curves represent axons and red dots represent terminals along axons. Purple triangles indicate en passant swellings (varicosity), while yellow triangles indicate terminals with necks. Dotted lines indicate linear distances between two adjacent axon terminals along the same axon, defined as terminal spacing. (E) Comparisons of distribution proportions of axonal varicosities from auditory L5 and L6 AT-projecting neurons across MGBv, MGBd, PIN and POL. Data were from 5 mice for L5 and 6 mice for L6. Field numbers for L5: 14 in MGBv, 9 in MGBd, PIN, and POL. Field numbers for L6: 18 in MGBv, 12 in MGBd, 12 in PIN, and 14 in POL. (F) Comparisons of distances between two adjacent axon terminals along the same axon from auditory L5 and L6 AT-projecting neurons across MGBv, MGBd, PIN and POL. Data were from 5 mice for L5 and 6 mice for L6. Fiber numbers for L5: 15 in MGBv, 12 in MGBd, 10 in PIN, and 10 in POL. Fiber numbers for L6: 22 in MGBv, 13 in MGBd, 13 in PIN, and 14 in POL. (G) Frequency distributions of sizes of axon terminals from rL5→AT and rL6→AT in the AT, including MGBv and MGBd, PIN and POL. Sample size of terminals: L5-MGBv (7505 terminals), MGBd (4964 terminals), MGBm (14520 terminals), PIN (9080 terminals), and POL (11239 terminals) from 5 mice; L6 - MGBv (92948 terminals), MGBd (44358 terminals), MGBm (561 terminals), PIN (2917 terminals), and POL (1041 terminals) from 6 mice. (H-I) Comparisons of terminal sizes (H) and terminal size variance (I) between axon terminals from L5 AT-projecting neurons and those from L6 AT-projecting neurons across subnuclei of AT including MGBv, MGBd, MGBd, PIN and POL. Data are presented as mean±SEM. (J) Illustration of differences in terminal density (horizontal axis), terminal size (vertical axis) and varicosity proportion (color bar) between rL5→AT and rL6→AT projections to PIN/PP and those to MGBv/d. The coordinates of center dots of ellipses represent means of differences in terminal density and terminal size between PIN/PP and MGBv/d, respectively. The half axis length of ellipses represents SEM of differences in terminal density and terminal size between PIN/PP and MGBv/d, respectively. The color of ellipses represents means of differences in varicosity proportion between PIN/PP and MGBv/d. Ordinary one-way ANOVA with Dunnett’s multiple comparisons tests were performed on comparisons within rL5 across AT subdivisions in E, H, and I, and comparisons within rL6 across AT subdivisions. Kruskal-Wallis test with Dunn’s multiple comparisons test was used for comparing within rL6 across AT subdivisions in E and within rL5 across AT subdivisions in F. Brown-Forsythe and Welch ANOVA tests with Tamhane’s T2 multiple comparisons test was performed on comparisons within rL6 across AT subdivisions in F. Friedman test with Dunn’s multiple comparisons test was performed on comparisons within rL6 across AT subdivisions in H, respectively. Multiple unpaired t tests with correction for using the Holm-Sidak method were used for comparing the same AT subdivisions between rL5 and rL6 in E-I. P values, n.s. *P*>0.05, * *P*<0.05, ** *P*<0.01, *** *P*<0.001, **** *P*<0.0001. More details in Table S2.

Based on the location of swellings along the axons, we classified axon terminals into two types, which are the axonal varicosities and boutons. Axonal varicosities are *en passant* presynaptic terminals that appear as small swellings of the axon (Fig 4C-D, indicated by purple triangles), and axonal boutons are terminal bulbs at the end of axons^75–78^ (Fig 4C-D, indicated by yellow triangles). In the MGB, about 70% of examined axon terminals from both L5 and L6 were varicosities (Fig 4E). In the PIN and POL, about 92% and 80% of L5 and L6 axon terminals were varicosities, respectively (Fig 4E). Therefore, varicosity was the dominating terminal type in all PoA subdivisions, and there are proportionally more axonal varicosities in the PIN and POL than in the MGB. Furthermore, the terminals along L5 and L6 axons showed sparser distribution in the PIN and POL compared with those in the MGB (Fig 3F, PIN/POL: 9.9±1.2 μm; MGB: 4.8±0.3 μm), and there was no significant difference in the spacing of L5 and L6 terminals in each individual AT subdivisions.

By reconstructing over a hundred thousand terminals and estimating terminal size with corrections (Fig S9; see Methods for a detailed description of correction procedure), we next conducted comparisons in the size of cortical axon terminals across the MGBv, MGBd, PIN and POL. In the MGBv/MGBd and PIN/POL, the distribution range of L5 terminal size was 0.4-2.0 μm and 0.4-2.8 μm with the size of 1.2 μm and 1.4 μm more commonly observed, respectively (Fig 4G, lines with filled circles). Regarding L6 terminals, the distribution range (0.4-2.0 μm) of terminal size was similar across AT subdivisions, and the size of 1.2 μm was more commonly observed (Fig 4G, lines with filled triangles). Thus, L5 terminals demonstrated a significantly larger variance in their size between the MGBv/MGBd and PIN/POL compared with L6 terminals (Fig 4H-I). In another word, the size of L6 terminals was more uniform across AT subdivisions. In addition, L5 terminals in the PIN/POL were significantly larger and more variant than L6 terminals, although they were similar in the MGBv/MGBd (Fig 4H-I).

From the characterization of axon terminal distribution and morphology features, we determined that rL5→AT and rL6→AT circuits are distinct types of corticothalamic pathways, which can be reflected by the differences between their innervation of AT subdivisions with distinct module selectivity (Fig 6J). rL5→AT terminals had the larger size, larger size variance and higher proportion of varicosity type in the PoA, while rL6→AT terminals showed larger density in the principal nucleus of the AT.

### Auditory thalamus-projecting L5 neurons are PT neurons

We next determined the cell type of rL5→AT neurons. First of all, our anti-GABA immunostaining data showed that rL5→AT neurons were not GABAergic (Fig 5A). We further confirmed this observation by injecting rAAV2-retro-FLEX-eGFP into the AT of PV-Cre, SST-Cre, or VIP-Cre mice (Fig S10A-B). As expected, no cortical neurons expressing eGFP were observed in the above three mouse lines (Fig S10C). Thus, rL5→AT neurons were exclusively excitatory. Furthermore, sparsely labeled rL5→AT neurons exhibited typical morphological characteristics of pyramidal neurons (Fig S10D-E), and rL5→AT neurons sent long-distance projections to the brainstem (Fig S10F), strongly suggesting that they are pyramidal tract (PT) neurons. Therefore, we conducted immunostaining for Ctip2(Fig 5B), COUP-TF-interacting protein 2, which has been considered as a molecular marker specific for PT neurons^79–81^, and found that about 99.4% of rL5→AT neurons examined were Ctip2 positive (Fig 5C), indicating that rL5→AT neurons were indeed PT neurons.

**Figure 5.**
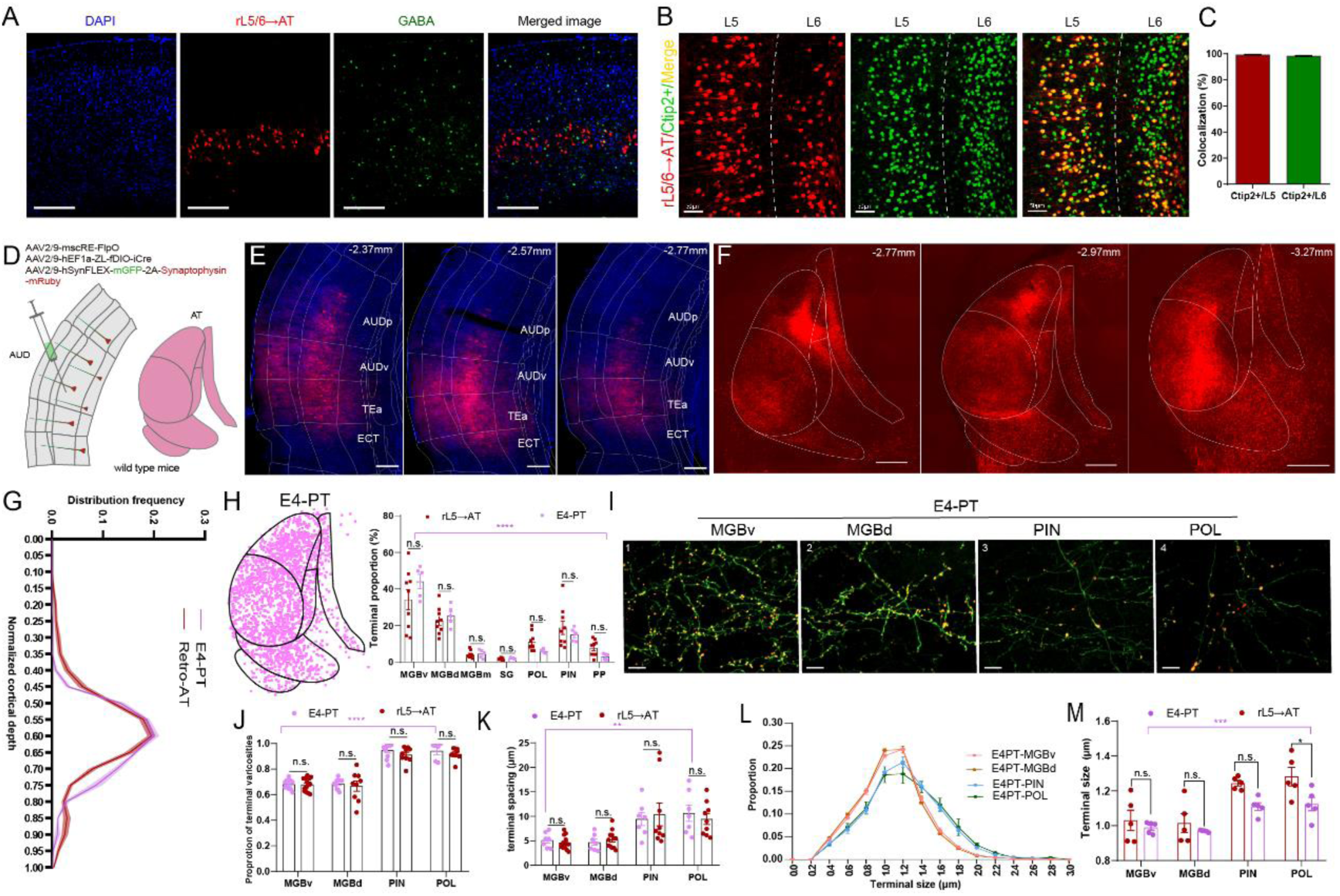
rL5→AT neurons are PT neurons with similarities in characteristics of soma distribution and corticothalamic circuits with E4-PT neurons. (A) Immunofluorescence staining of GABA (Green) in brain slices containing retrogradely labeled AT-projecting neurons (Red). Scale bars, 200um. (B) Immunofluorescent staining of Ctip2 in the auditory cortex showing retrogradely labeled L5 and L6 AT- projecting neurons. Green, Ctip2 positive neurons. Red, AT-projecting neurons expressing tdTomato. Orange, colocalizations of AT-projecting neurons with Ctip2 positive neurons. Scale bars, 50μm. (C) Proportions of rAAV2-retro-labeled AT-projecting neurons expressing Ctip2. Data are presented as mean±SEM and were collected from 10 slices of 2 mice. (D) Schematic of fluorescent-labeled auditory cortical pyramidal tract (PT) neurons and their axon terminals, combining viral tools with mscRE enhancer and double recombinase systems. (E) Confocal images of PT neurons expressing synaptophysin-mRuby specifically driven by mscRE enhancer in the auditory cortices. Blue, DAPI. Red, synaptophysin-mRuby. Scale bars, 200μm. (F) Confocal images illustrating axon terminals of PT neurons in the auditory thalamus at different AP distances. Scale bars, 200μm. (G) Illustration of distribution frequencies of E4-PT neurons and AT-projecting neurons along the normalized depth of auditory cortices. (H) Left, three thousand points which were distributed in the auditory thalamus representing axon terminals of PT neurons from weighted samples. Right, Comparisons of axon terminal distribution in the subnuclei of the auditory thalamus between L5 AT-projecting neurons retrogradely labeled by rAAV2-retro (n=5) and PT neurons specifically labeled by PT enhancer (n=5). Data are presented as mean±SEM. (I) Representative confocal images displaying axons (green) and terminals (overlap of green and red) distributed in MGBv (I1), MGBd (I2), PIN (I3), and POL (I4) from E4-PT neurons. Scale bars, 10μm. (J) Comparisons of proportions of varicosity-type between terminals from E4-PT neurons and those from rL5→AT neurons to the MGBv, MGBd, PIN and POL. Data samples for rL5→AT are the same in the Figure 4. Data samples for E4-PT were from 5 mice. Field numbers for E4, 11 in MGBv, 11 in MGBd, 10 in PIN, and 6 in POL. Fiber numbers for E4: 7 in MGBv, 7 in MGBd, 7 in PIN, and 7 in POL. (K) Comparison of terminal spacing between axon terminals from E4-PT neurons and those from rL5→AT neurons to subnuclei of AT including MGBv, MGBd, PIN and POL. Data are presented as mean±SEM. (L) Frequency distributions of axon terminal sizes from E4-PT neurons to MGB including MGBv and MGBd, as well as PIN and POL. Sample sizes: MGBv (24753 terminals), MGBd (12938), PIN (9656), and POL (1581) from 5 mice. (M) Comparison of terminal sizes between axon terminals from E4-PT neurons and those from rL5→AT neurons to subnuclei of AT including MGBv, MGBd, PIN and POL. Data are presented as mean±SEM. Multiple unpaired t tests with correction for using the Holm-Sidak method were used for comparisons in the same AT subdivisions between E4-PT and rL5→AT. For comparisons within E4-PT or rL5→AT across AT subdivisions, Kruskal-Wallis test with Dunn’s multiple comparisons test, ordinary one-way ANOVA with Tukey’s multiple comparisons test, and RM one-way ANOVA with Tukey’s multiple comparisons test were used in H, K, and M. P values, n.s. *P*>0.05, ** *P*<0.01, **** *P*<0.0001. More details in Table S2.

Since the rL5→AT neurons may only represent a subpopulation of PT neurons in the cortex, would the distribution pattern and morphological features of the axon terminals of these neurons be representative those of PT neurons’ axon terminals in the AT? By injecting mscRE4 enhancer virus^47^ into the deep layers of the AUD, we specifically labeled the soma and axon terminals of PT neurons using Synaptophysin-mRuby (Fig 5D-F, E4-PT). We found that, first, E4-PT neurons were mainly located in middle L5 (Fig 5G), resembling the distribution of rL5→AT neurons. Second, similar to the distribution pattern of rL5→AT terminals (Fig 5H), dense E4-PT axon terminals were observed throughout the entire AT (Fig 5F&H). Third, the morphological features of E4-PT axon terminals (Fig 5I), including terminal type (Fig 5J) and spacing (Fig 5K), closely resembled those of rL5→AT axon terminals, although the size of them tended to be smaller in the association module of the AT (Fig 5L-M), suggesting that rL5→AT neurons are a subset of E4-PT neurons with larger terminal size. Taken together, these data indicated that rL5→AT circuit is indeed a typical PT-originated corticothalamic pathway.

We also examined the cell type of rL6→AT neurons, although they only represented a minor group of neurons labeled by rAAV2-retro. We found that these neurons were excitatory neurons as well (Fig 5A), and that, interestingly, about 98.4% of these neurons were Ctip2 positive (Fig 5C). Nevertheless, 97.3% of rL6→AT neurons were also immunopositive for Tbr1 (Fig S11A-B), T-box brain transcription factor1, which is a molecular marker for L6 corticothalamic (CT) neurons^79,82^ and also expressed by RL6→AT neurons (Fig S11C). This high rate of co-expression indicated that rL6→AT neurons are a special group of L6 neurons. Considering that RL6→AT neurons greatly outnumbered rL6→AT neurons, that both groups are CT neurons, and that they showed similar laminar distribution patterns, rL6→AT neurons were very likely a subpopulation of RL6→AT neurons.

We wondered whether rL6→AT neurons would show characteristics of CT neurons in terms of connectivity. Toward this end, we specifically labeled the soma and axon terminals of CT neurons by injecting AAV9-Flex-mGFP-2A-Synaptophysin-mRuby into the AUD of Ntsr1-Cre mouse line (Fig S11D-F, Ntsr1+), which has been widely used to investigate L6 CT circuits^70–73^. Similar to rL6→AT neurons, Ntsr1+ neurons were predominantly distributed in middle L6 (Fig S11G). In addition, the distribution pattern (Fig S10F), type (Fig S11J) and spacing (Fig S11K) of Ntsr1+ terminals resembled those of rL6→AT neurons as well. Nevertheless, it should be noted that rL6→AT terminals in the primary and non-primary modules of the AT tended to be larger than Ntsr1+ terminals (Fig S11L-M). These data collectively indicated that rL6→AT neurons are a subset of CT neurons with larger terminals.

### Fully connected macroscale architecture of L5-originated corticothalamic pathways

Taken together, we propose a novel architecture for the corticothalamic connectivity originating from L5 PT neurons in the auditory system. In this architecture, the temporal association cortex (TEa) contributes the most L5 PT neurons projecting to each individual module, including the primary module of the AT, followed by the secondary auditory cortices (AUDv and AUDd) and primary auditory cortex (AUDp) (Fig 6A and left panel in B). In the AT, axon terminals from L5 PT neurons are nearly evenly distributed across different modules, although those in the association module are significantly larger than those in the primary and non-primary modules (Fig 6B middle and right panels). Thus, in the auditory system, the L5→thalamus pathways form a projection pattern that resembles a fully connected neural network^83–85^ at macroscale. This fully-connected-like architecture proposed here is distinct from the traditional one that emphasizes hierarchical projections from L5 neurons residing in lower-order sensory cortices to higher-order sensory thalamic nuclei^35,86–88^.

**Figure 6.**
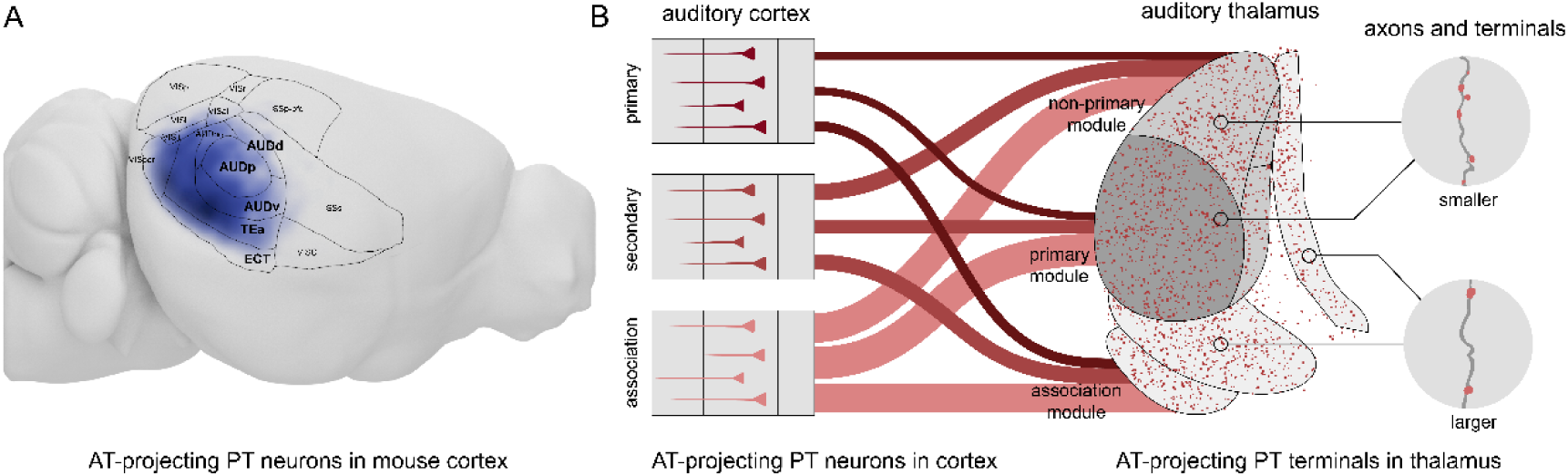
Fully-connected architecture of L5 PT-originated corticothalamic pathways. (A) Relative density estimation of AT-projecting PT neurons in the temporal cortex of mice. The darker the blue, the higher density in the location. (B) Connectivity between different cortical areas, including the primary and secondary auditory cortex and temporal association cortex, and different modules of the auditory thalamus. Three lines in different colors to each individual module of the auditory thalamus and their widths represent proportions of L5 PT neurons projecting to the AT module from the cortical area accounting for those from the entire cortical areas. In the middle panel, red dots represent axon terminal distributed in the AT. In the right panel, cartooned axon with smaller terminals distributed on it indicate axon and terminals from L5 PT neurons to the primary and non-primary modules of the auditory thalamus, and the cartooned axon with larger terminals indicate those distributed in the polymodal association AT module.

## Discussion

It has long been believed that corticothalamic projections in the sensory systems play a critical role in enabling adaptive information processing and transmission in the thalamus. However, the data that can be used to enlighten the organizing principles of the projections from L5 to the thalamus remained far from sufficient. This fact is out of balance with the widely recognized functional importance of the long-range projections of L5 neurons. Here, by using rAAV2-retro and various cutting-edge circuit-mapping techniques, we found that each individual subdivision of the auditory thalamus is innervated by a large number of L5 neurons, which are most numerous in the TEa followed by the secondary and primary auditory cortices. These findings provide a new anatomical framework for functionally interrogating the corticothalamic circuits originating from the L5 of sensory cortices, which may facilitate the emergence of a substantially revised corticothalamic model in the sensory systems.

### Strong L5 projections to the primary thalamic nucleus across sensory modalities

The preference of rAAV2-retro for cortical L5 neurons has been demonstrated by several studies injecting this viral tracer into L5-targetted subcortical structures^9,13,46,72,89–91^. By using rAAV2-retro, we showed that a large number of L5 neurons in the primary auditory, visual and somatosensory cortices make direct projections to their corresponding primary sensory thalamic nuclei. This finding is consistent with that reported by a recent study using the same virus in the visual system^92^. We further found that, to our surprise, the secondary sensory and temporal association cortices send L5 projections to the primary sensory thalamic nuclei as well, and that the TEa is the dominating provider of L5 projections. Thus, the circuit model suggested by our data is distinct from the long-proposed lower order L5→higher order thalamus model^35,86–88^. The influence of L5 neurons on sensory processing in the primary thalamic nuclei may be much more complicated and powerful than previously thought.

### The TEa is a major source of cortical information for the auditory thalamus

Consistent with previous observations^93^, MGBv-projecting L6 CT neurons were concentrated in the AUDp, indicating that the MGBv receives feedback information from the L6. However, L6 CT neurons that project to the MGBd and PoA were mainly found in the TEa followed by the AUDv. Thus, the distribution patterns of L6 CT neurons are distinct between the primary and non-primary/polymodal association modules. Surprisingly, the distribution patterns of L5 neurons projecting to different AT modules are quite uniform. Specifically, the TEa provided the most L5 neurons for each individual module of the AT, followed by the AUDv and AUDp, suggesting that descending cortical information from L5 may not be relevant to the specific function of an AT module in auditory processing. Collectively, our data showed that the TEa turns out to be a major source of cortical information for the entire AT. As one of the central hubs in the cortex^94^, the TEa may provide the AT with highly processed sensory information and contextual information, which includes both external needs^95^ and internal states^96^, to shape thalamic auditory processing, contributing to a coherent and adaptive representation of the sensory world.

### L5 and L6 neurons use complementary strategies to preferentially innervate distinct AT modules

At the population level, both the size and density of axon terminals have been used to infer the effectiveness of presynaptic inputs in changing postsynaptic membrane potentials^97–104^. Larger and denser axon terminals may have a greater impact on postsynaptic neurons^105,106^. Interestingly, we found that the average size of L5 terminals in the association module is significantly larger than that in the other two modules, although the density was similar. In contrast, L6 terminals are significantly denser in the primary and non-primary modules than in the association module, but their average size did not correlate with specific AT modules. Given that AT-projecting L5 neurons receive significantly more inputs from the amygdala and striatum whereas L6 CT neurons integrate more inputs of multiple modalities (Fig S12), L5 and L6 neurons may use size- and density-dominated strategies to preferentially relay contextual information and processed sensory information to the association and primary/non-primary modules, respectively.

### Potential technical limitation

In the present study, we showed that a large number of cortical PT neurons were retrogradely labeled when rAAV2-retro was injected into the auditory thalamus. In fact, it has been widely acknowledged that different tracers and viral tools for retrograde tracing prefer to label different types of presynaptic neurons, especially for cortical neurons^46,107,72,89–92^. Therefore, it is possible rAAV2-retro only labeled a subset of PT neurons projecting to the auditory thalamus. Nevertheless, the laminar distribution and projection patterns of PT neurons labeled by the mscRE4 virus, which selectively targets and labels cortical PT neurons, resembled those labeled by rAAV2-retro, suggesting that rAAV2-retro-labeled PT neurons represent a major subset of AT- projecting PT neurons (Figure 5). According to these data, the PT→thalamus circuit framework derived from rAAV2-retro labeling may still hold even if novel subtypes of PT neurons are identified in the future.

## Acknowledgments

We thank Y. Yue for immumohistochemical staining support, B. Hong for instrumental support, M. M. Luo and Z. C. Guo for mice and viruses supplement, and X. H. Zhang and Y. Q. Li for critical comments on the manuscript.

K.Y. receives funding from STI 2030-Major Projects 2021ZD0200300, National Natural Science Foundation of China (31871057, 32070993, 81527901, T2341003), Beijing Municipal Science & Technology Commission (Z181100001518004, Z181100001518006), and Guoqiang Institute, Tsinghua University.

## Author contributions

Conceptualization, K.Y. and F.X.; Methodology and Investigation, F.X., Y.G., T.W. and M.L.; Data analysis, F.X., Y.G. and T.W.; Data Curation, all authors; Writing – Original Draft, K.Y., F.X. and Y.G.; Writing – Review & Editing, K.Y., F.X. and Y.G.; Supervision, K.Y.

## STAR⋆METHODS

### KEY RESOURCES TABLE

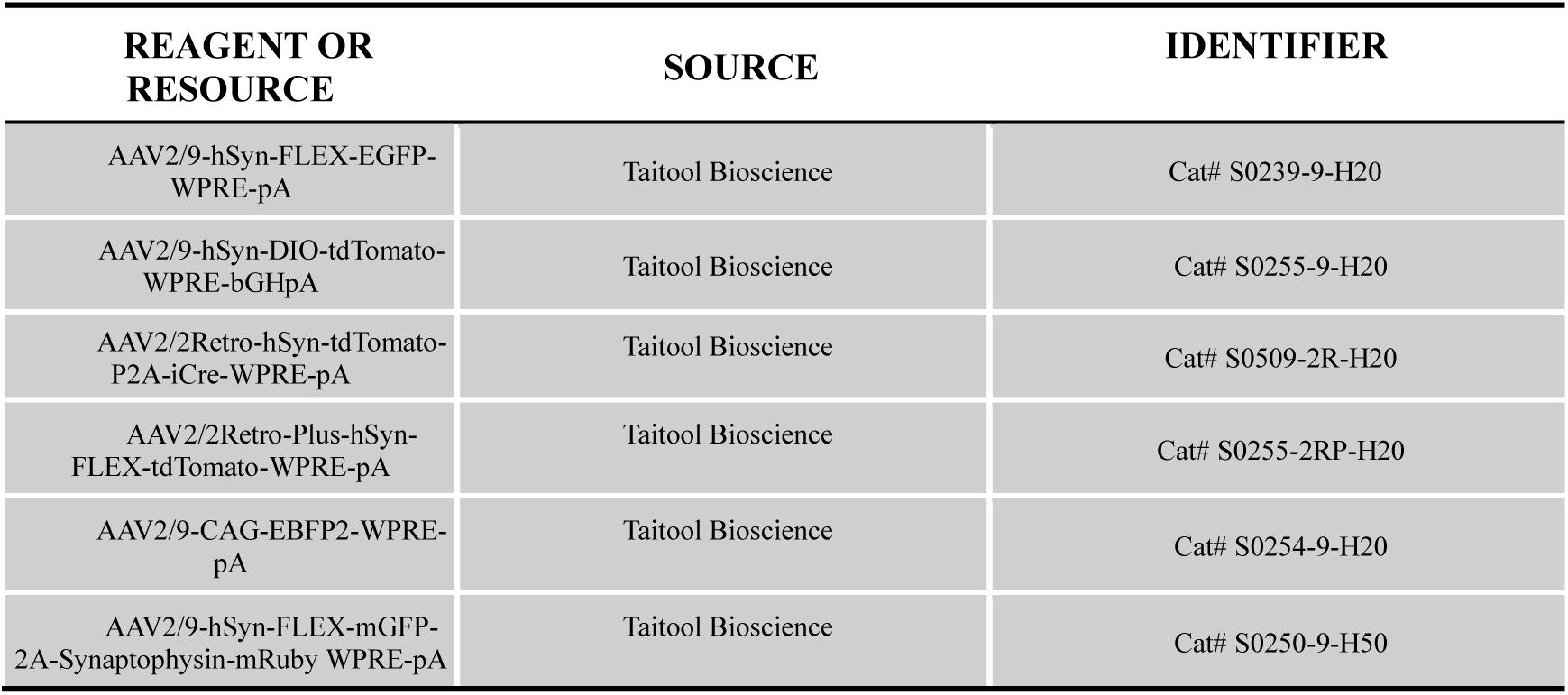
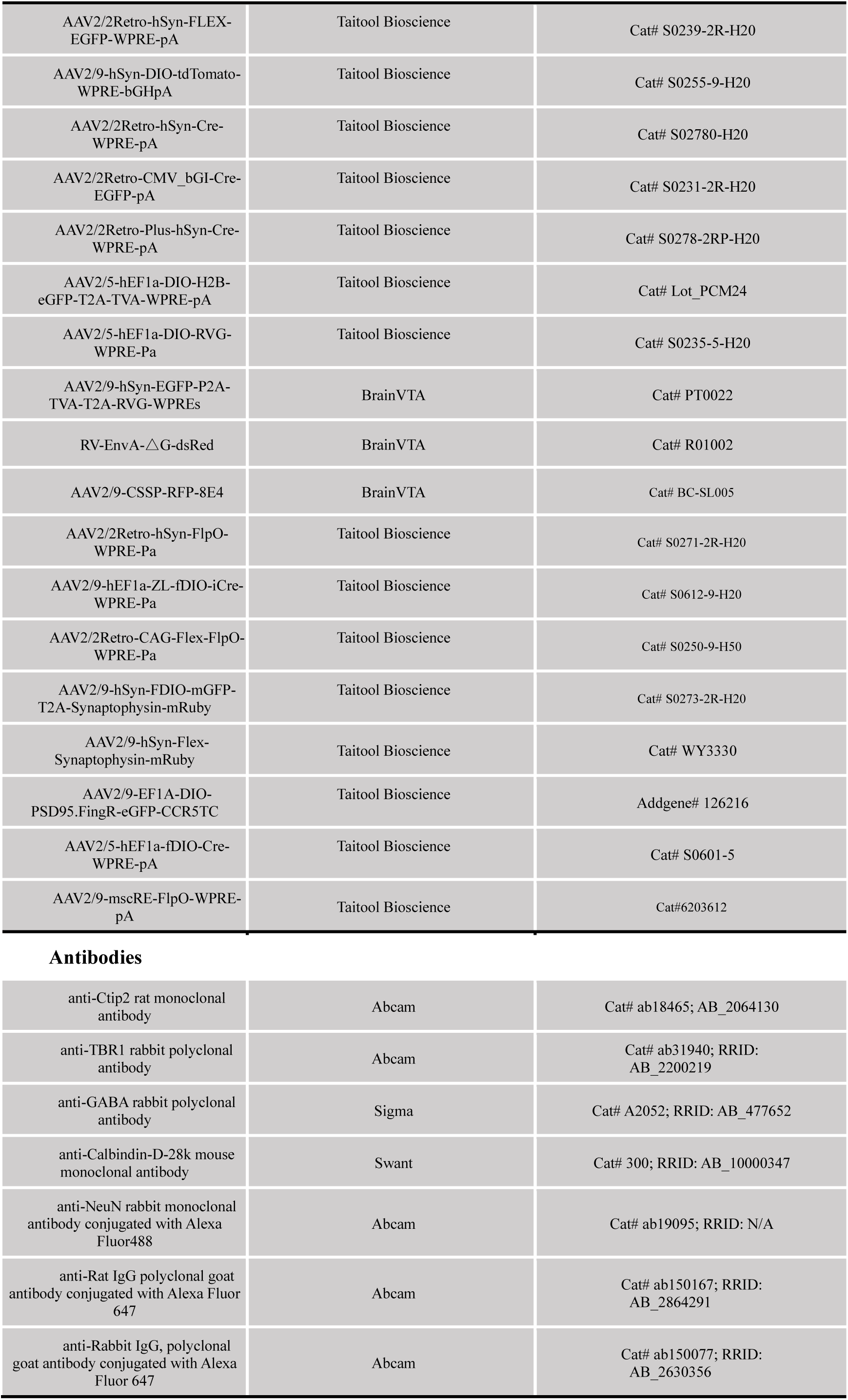
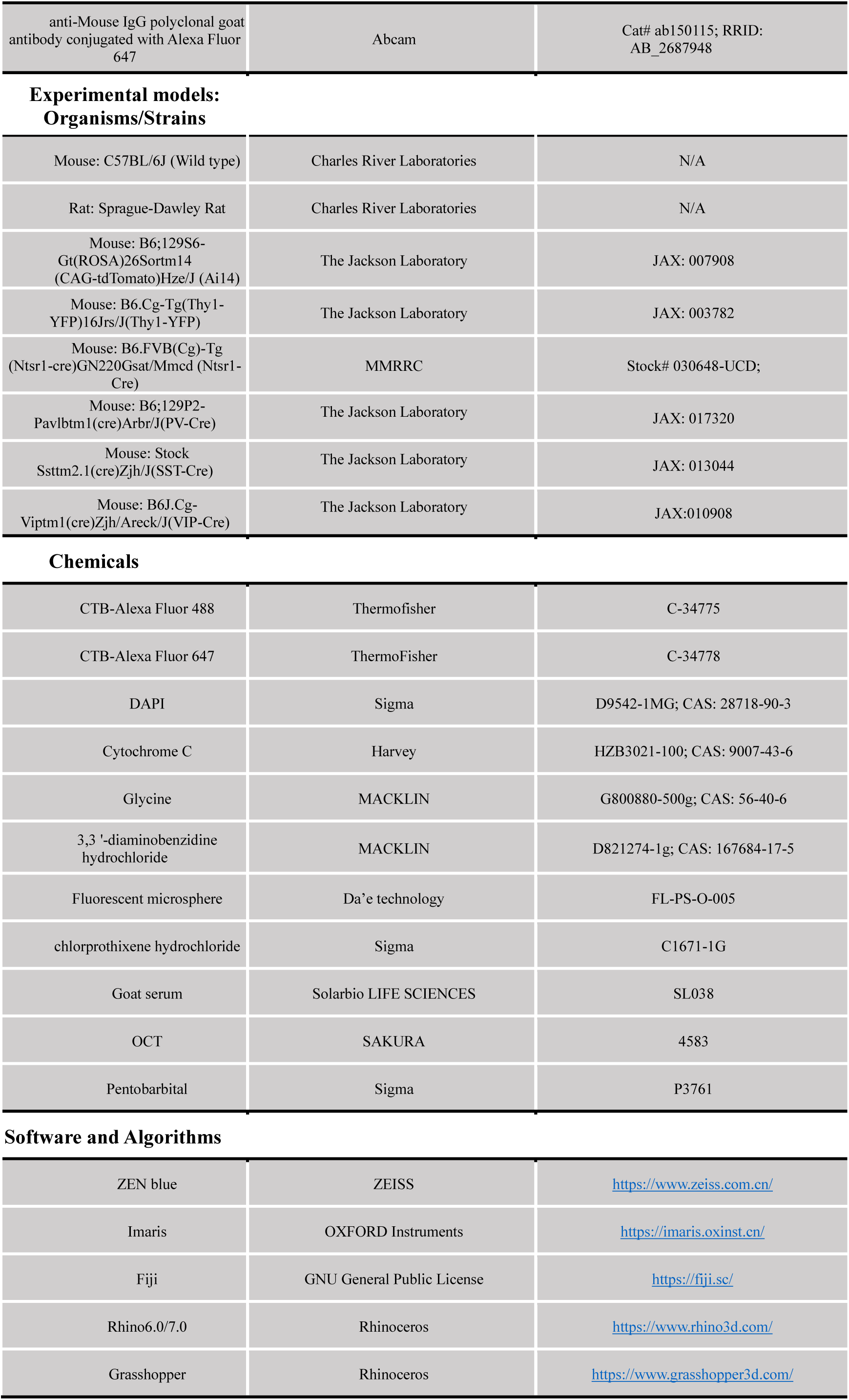
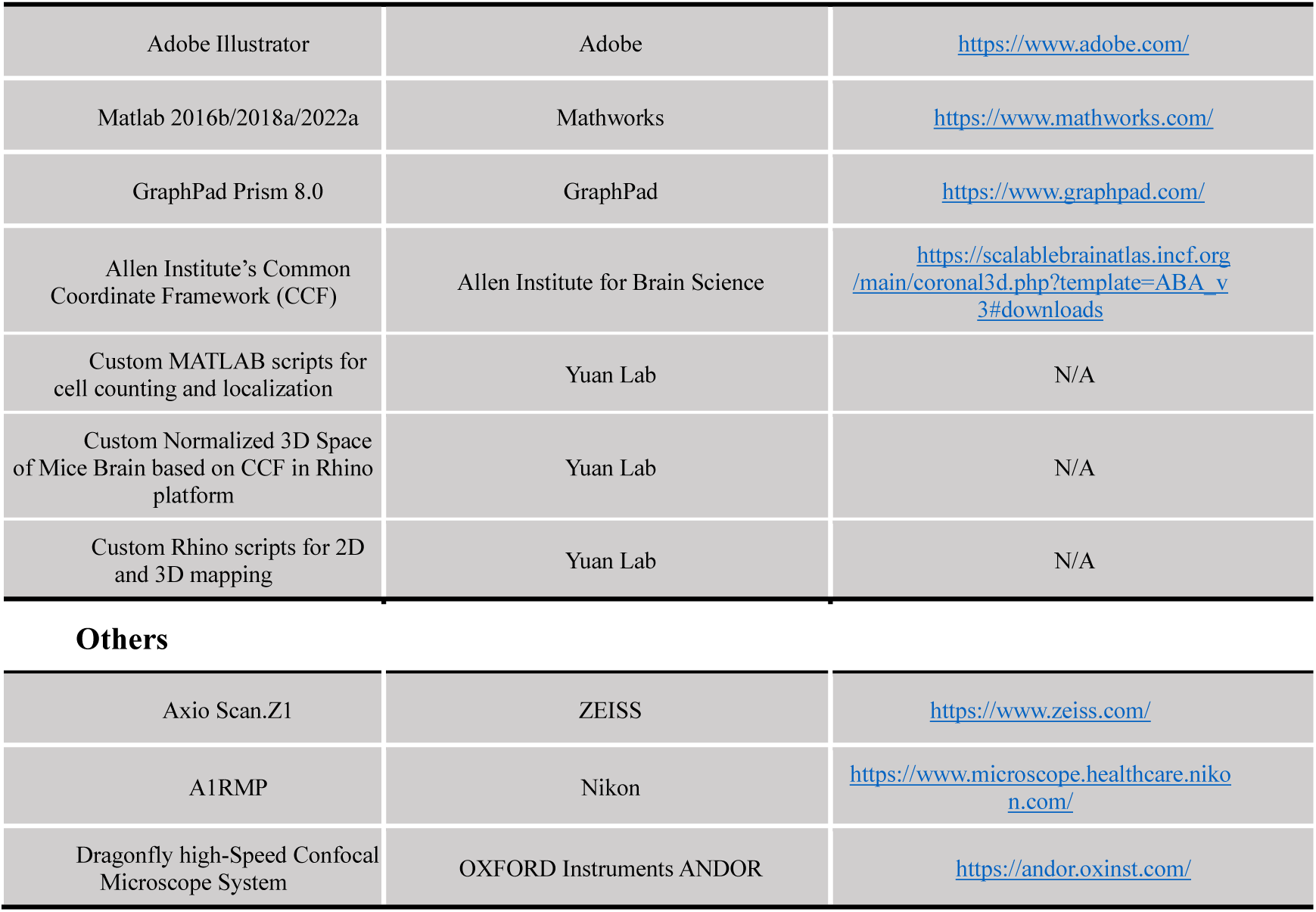

| REAGENT OR RESOURCE | SOURCE | IDENTIFIER |
| --- | --- | --- |
| AAV2/9-hSyn-FLEX-EGFP-WPRE-pA | Taitool Bioscience | Cat# S0239-9-H20 |
| AAV2/9-hSyn-DIO-tdTomato-WPRE-bGHpA | Taitool Bioscience | Cat# S0255-9-H20 |
| AAV2/2Retro-hSyn-tdTomato-P2A-iCre-WPRE-pA | Taitool Bioscience | Cat# S0509-2R-H20 |
| AAV2/2Retro-Plus-hSyn-FLEX-tdTomato-WPRE-pA | Taitool Bioscience | Cat# S0255-2RP-H20 |
| AAV2/9-CAG-EBFP2-WPRE-pA | Taitool Bioscience | Cat# S0254-9-H20 |
| AAV2/9-hSyn-FLEX-mGFP-2A-Synaptophysin-mRuby WPRE-pA | Taitool Bioscience | Cat# S0250-9-H50 |
| AAV2/2Retro-hSyn-FLEX-EGFP-WPRE-pA | Taitool Bioscience | Cat# S0239-2R-H20 |
| AAV2/9-hSyn-DIO-tdTomato-WPRE-bGHpA | Taitool Bioscience | Cat# S0255-9-H20 |
| AAV2/2Retro-hSyn-Cre-WPRE-pA | Taitool Bioscience | Cat# S02780-H20 |
| AAV2/2Retro-CMV_bGII-Cre-EGFP-pA | Taitool Bioscience | Cat# S0231-2R-H20 |
| AAV2/2Retro-Plus-hSyn-Cre-WPRE-pA | Taitool Bioscience | Cat# S0278-2RP-H20 |
| AAV2/5-hEF1a-DIO-H2B-eGFP-T2A-TVA-WPRE-pA | Taitool Bioscience | Cat# Lot_PCM24 |
| AAV2/5-hEF1a-DIO-RVG-WPRE-Pa | Taitool Bioscience | Cat# S0235-5-H20 |
| AAV2/9-hSyn-EGFP-P2A-TVA-T2A-RVG-WPREs | BrainVTA | Cat# PT0022 |
| RV-EnvA-△G-dsRed | BrainVTA | Cat# R01002 |
| AAV2/9-CSSP-RFP-8E4 | BrainVTA | Cat# BC-SL005 |
| AAV2/2Retro-hSyn-FlpO-WPRE-Pa | Taitool Bioscience | Cat# S0271-2R-H20 |
| AAV2/9-hEF1a-ZL-fDIO-iCre-WPRE-Pa | Taitool Bioscience | Cat# S0612-9-H20 |
| AAV2/2Retro-CAG-Flex-FlpO-WPRE-Pa | Taitool Bioscience | Cat# S0250-9-H50 |
| AAV2/9-hSyn-FDIO-mGFP-T2A-Synaptophysin-mRuby | Taitool Bioscience | Cat# S0273-2R-H20 |
| AAV2/9-hSyn-Flex-Synaptophysin-mRuby | Taitool Bioscience | Cat# WY3330 |
| AAV2/9-EF1A-DIO-PSD95.FingR-eGFP-CCR5TC | Taitool Bioscience | Addgene# 126216 |
| AAV2/5-hEF1a-fDIO-Cre-WPRE-pA | Taitool Bioscience | Cat# S0601-5 |
| AAV2/9-mscRE-FlpO-WPRE-pA | Taitool Bioscience | Cat#6203612 |
| Antibodies |  |  |
| anti-Ctip2 rat monoclonal antibody | Abcam | Cat# ab18465; AB_2064130 |
| anti-TBR1 rabbit polyclonal antibody | Abcam | Cat# ab31940; RRID: AB_2200219 |
| anti-GABA rabbit polyclonal antibody | Sigma | Cat# A2052; RRID: AB_477652 |
| anti-Calbindin-D-28k mouse monoclonal antibody | Swant | Cat# 300; RRID: AB_10000347 |
| anti-NeuN rabbit monoclonal antibody conjugated with Alexa Fluor488 | Abcam | Cat# ab19095; RRID: N/A |
| anti-Rat IgG polyclonal goat antibody conjugated with Alexa Fluor 647 | Abcam | Cat# ab150167; RRID: AB_2864291 |
| anti-Rabbit IgG, polyclonal goat antibody conjugated with Alexa Fluor 647 | Abcam | Cat# ab150077; RRID: AB_2630356 |
| anti-Mouse IgG polyclonal goat antibody conjugated with Alexa Fluor 647 | Abcam | Cat# ab150115; RRID: AB_2687948 |
| Experimental models: Organisms/Strains |  |  |
| Mouse: C57BL/6J (Wild type) | Charles River Laboratories | N/A |
| Rat: Sprague-Dawley Rat | Charles River Laboratories | N/A |
| Mouse: B6.129S6-Gt(ROSA)26Sortm14 (CAG-tdTomato)Hze/J (Ai14) | The Jackson Laboratory | JAX: 007908 |
| Mouse: B6.Cg-Tg(Thy1-YFP)16Jrs/J(Thy1-YFP) | The Jackson Laboratory | JAX: 003782 |
| Mouse: B6.FVB(Cg)-Tg (Ntsr1-cre)GN220Gsat/Mmcd (Ntsr1-Cre) | MMRRC | Stock# 030648-UCD; |
| Mouse: B6.129P2-Pavlbtm1(cre)Arbr/J(PV-Cre) | The Jackson Laboratory | JAX: 017320 |
| Mouse: Stock Ssttm2.1(cre)Zjh/J(SST-Cre) | The Jackson Laboratory | JAX: 013044 |
| Mouse: B6J.Cg-Viptm1(cre)Zjh/Arck/J(VIP-Cre) | The Jackson Laboratory | JAX:010908 |
| Chemicals |  |  |
| CTB-Alexa Fluor 488 | ThermoFisher | C-34775 |
| CTB-Alexa Fluor 647 | ThermoFisher | C-34778 |
| DAPI | Sigma | D9542-1MG; CAS: 28718-90-3 |
| Cytochrome C | Harvey | HZB3021-100; CAS: 9007-43-6 |
| Glycine | MACKLIN | G800880-500g; CAS: 56-40-6 |
| 3,3 '-diaminobenzidine hydrochloride | MACKLIN | D821274-1g; CAS: 167684-17-5 |
| Fluorescent microsphere | Da'e technology | FL-PS-O-005 |
| chlorprothixene hydrochloride | Sigma | C1671-1G |
| Goat serum | Solarbio LIFE SCIENCES | SL038 |
| OCT | SAKURA | 4583 |
| Pentobarbital | Sigma | P3761 |
| Software and Algorithms |  |  |
| ZEN blue | ZEISS | https://www.zeiss.com.cn/ |
| Imaris | OXFORD Instruments | https://imaris.oxinst.cn/ |
| Fiji | GNU General Public License | https://fiji.sc/ |
| Rhino6.0/7.0 | Rhinoceros | https://www.rhino3d.com/ |
| Grasshopper | Rhinoceros | https://www.grasshopper3d.com/ |
| Adobe Illustrator | Adobe | https://www.adobe.com/ |
| Matlab 2016b/2018a/2022a | Mathworks | https://www.mathworks.com/ |
| GraphPad Prism 8.0 | GraphPad | https://www.graphpad.com/ |
| Allen Institute's Common Coordinate Framework (CCF) | Allen Institute for Brain Science | https://scalablebrainatlas.incf.org/main/coronal3d.php?template=ABA_v3#downloads |
| Custom MATLAB scripts for cell counting and localization | Yuan Lab | N/A |
| Custom Normalized 3D Space of Mice Brain based on CCF in Rhino platform | Yuan Lab | N/A |
| Custom Rhino scripts for 2D and 3D mapping | Yuan Lab | N/A |
| Others |  |  |
| Axio Scan.Z1 | ZEISS | https://www.zeiss.com/ |
| A1RMP | Nikon | https://www.microscope.healthcare.nikon.com/ |
| Dragonfly high-Speed Confocal Microscope System | OXFORD Instruments ANDOR | https://andor.oxinst.com/ |

### CONTACT FOR REAGENT AND RESOURCE SHARING

Further information and requests for resources and reagents should be directed to and will be filled by the Lead Contact, Kexin Yuan.

### EXPERIMENTAL MODEL AND SUBJECT DETAILS

#### Animals

All animal care procedures and experiments were approved by the Institutional Animal Care and Use Committee at Tsinghua University, Beijing, China. Both before and after surgery, all the mice were housed with standard temperature, humidity, and light/dark cycle conditions in the Laboratory Animal Resources Center, Tsinghua University. Adult mice (2-4 months, Background strain C57B6/J) of both sexes were used. Wild-type mice and rats were purchased from Wei Tong Li Hua Experimental Animal Co., Ltd (Beijing China). The transgenic mice, Ai14 (B6. Cg-Gt (ROSA)26Sortm14(CAG-tdTomato)Hze/J, Stock #: 007914), from the Jackson laboratories. The Thy1-YFP-H^49,50^ mice (B6. Cg-Tg(Thy1-YFP)16Jrs/J, Stock #: 003782) were used for histochemistry parcellation of auditory thalamus. The Ntsr1-Cre (B6.FVB(Cg)-Tg (Ntsr1-cre) GN220Gsat/Mmcd, Stock #: 030648-UCD) mice from the MMRRC were used for labeling L6 neurons, respectively. The PV-Cre (B6;129P2-Pavlbtm1(cre)Arbr/J, Stock #: 017320), SST-Cre (Stock Ssttm2.1(cre)Zjh/J, Stock #: 013044) and Vip-Cre (B6J.Cg-Viptm1(cre)Zjh/Areck/J, Stock#: 010908) mice from the Jackson Lab were all used for labeling the cortical inhibitory neurons.

### EXPERIMENTAL DETAILS

#### Abbreviations for brain regions

Most brain regions abbreviation were from the Allen Mouse Brain Atlas^54^ (https://scalablebrainatlas.incf.org/main/coronal3d.php?template=ABA_v3#downloads) and the Mouse Brain in Stereotaxic Coordinates, second edition, Franklin, K. B. J. and Paxinos, G ^108^.

#### Stereotaxic surgeries

To begin the experiments, mice were anesthetized with pentobarbital (i.p. 100mg/kg) and secured in a stereotaxic apparatus (RWD, 68001, Shenzhen, China). Erythromycin eye ointment was applied to prevent eye drying and an electric heating pad was used to maintain the body temperature of mice throughout the surgery process. Using sterilized scissors, a small incision was made in the skin over the skull midline and forceps were used to expose the bregma, lambda and skull surface. A small craniotomy was then performed above the auditory cortex and the auditory thalamus. Viruses were slowly injected into the bilateral auditory cortex and the auditory thalamus using a micro syringe pump with the glass pipette (4878, WPI, USA) at a rate of 50-100nl/min respectively. The following coordinates (in mm) were used ^108^: -2.70 AP, ±4.30 ML,-0.80/-1.20/-1.50 DV for the auditory cortex; -3.00 AP, ±2.10 ML, -3.00 DV for MGBv (AP is relative to bregma; ML is relative to the midline; DV is relative to the brain tissue surface above the targeted brain areas). The glass pipette was left in place for another 2-5 min after injection and then slowly withdrawn. After retracting the pipette, the skin was sutured. Following the surgery, mice were allowed to recover on a heated pad until ambulatory and then returned to their home cage. Mice underwent a post-surgery recovery period of approximately 3-4 weeks to allow for AAV virus infection and gene expression.

#### Virus and tracer injection

Viruses with the corresponding volumes and titers were injected and injection timelines were listed in the Table S1

#### Histology and Immunochemistry

Animals were administered an overdose of pentobarbital sodium salt (i.p.,300mg/kg) or avertin (i.p., 2.5%, 0.4ml/mice) and then underwent transcardial perfusion with saline (0.9%) followed by 4% paraformaldehyde (PFA) in the 0.01M phosphate-buffered saline (PBS). Brains were dissected, post-fixed overnight in 4℃ using 4% PFA and then dehydrated and cryoprotected in 30% sucrose until they sank. Brains were then frozen in optimal cutting temperature compound (OCT, Sakura Tokyo, Japan) and 50μm coronal slices were obtained covering the whole brain using a freezing microtome (CM1950, Leica Biosystems, Germany). For brain-wide cell counting, every other slice was collected and mounted on gelatin-coated slides. For antibody immunofluorescent staining, 40- 50μm coronal brain sections were collected into the ultra-low attachment multiple well plates (Corning^®^ Costar^®^) filled with 0.01M PBS. Then the collected brain sections were rinsed 3 times for 5min each time with 1XPBS buffer on a shaker to wash off any remaining frozen section embedding agent and other impurities. To minimize the influence of PFA fixation on antigen-antibody binding, antigen repairing was optionally performed to expose the antigenic determinants that are sensitive to PFA. Brain slices were soaked in the antigen repair solution (P0090, Beyotine) for 5min and then rinsed several times (e.g., 3 times, 5mins for each time) to remove any residual repair solution. Next, brain sections were incubated for 3x15min with 0.3% Triton 100 in 0.01M PBS for permeabilization and incubated for 60 minutes with 5% goat serum (SL038, Solarbio) to block unspecific binding of the antibodies. Next, brain sections were incubated with primary antibodies overnight at 4℃, followed by 3X15 minutes washes with PBS and incubated with the secondary antibodies for 2 hours. Sections were further washed thoroughly with PBS and coverslipped with 50% glycerol mounting medium. Additionally, sections were stained with 1:5000 DAPI (D1306, Invitrogen, USA) in 50% glycerol to visualize nuclei. The primary antibodies used in the study were anti-CB-D28k mouse monoclonal antibody (1:1000, 300, Swant, Switzerland), anti-GABA rabbit polyclonal antibody (1:1000, A2052, Sigma), anti-Ctip2 rat monoclonal antibody (1:500, ab18465, Abcam), anti-TBR1 rabbit polyclonal (1:1000, ab31940, Abcam), and the secondary antibodies were anti-Rat IgG Goat antibody conjugated with Alexa Fluor 647 (1:200, ab150167, Abcam), anti-Rabbit IgG polyclonal goat antibody conjugated with Alexa Fluor 647 (1:200, ab150077, Abcam), anti-Mouse polyclonal goad antibody conjugated with Alexa Fluor 647 (1:200, ab150115, Abcam). For NeuN staining, anti-NeuN rabbit monoclonal antibody conjugated with Alexa Fluor488 (1:1000, ab19095, Abcam) were used without secondary antibody incubation. For cytochrome oxidase staining (CYO staining), we followed the methods described in the previous literature^109^. Brain sections containing the auditory thalamus, ranging from AP -2.57mm to AP -3.27mm, were collected and mounted on gelatin-coated slides. Solution 1# was prepared by dissolving 20mg 3 ’3-diamindobenzidine hydrochloride (DAB, D821274, MACKLIN) in 10ml ddH2O and then stored frozen at -20℃ before use. Solution 2# was prepared by dissolving 30mg cytochrome C (HZB3021-100, Harvey) and 3g sucrose in 30ml 0.01M PBS and stored at 4 ℃ before use. The mixture of solution 1# and solution 2# in a 1:3 ratio was prepared for staining. The brain sections on the slide placed in the sealed immunohistochemistry wet box were fully immersed in the staining solution at 37 ℃ for 3 to 5 hours.

#### Rapid and non-Scaling on-Slide Tissue Clearing

In the present study, we developed our original rapid and non-scaling on-slide tissue clearing method and combined it with the high-speed spinning-disk confocal imaging to visualize the microscale structure of corticothalamic neurons with high throughput and high precision. Firstly, mice were perfused transcardially with heparin-sodium chloride solution (11566427, MACKLIN) at a rate of 22-24ml/min for 2 minutes, followed by 4% PFA at the rate of 16ml/min for 10 minutes, both at 4℃. After the descending aortic branch was clipped, mice were perfused with a room-temperature equilibrium fixative which contained 4% PFA, 1% sodium deoxycholate SDC (C12156670, MACKLIN) and 0.3M trehalose (S28J12H138585, Yuanye Bio-Technology) in ddH2O at a rate of 4ml/min for 10 minutes. Then dissected brains were immersed in a balanced cryopreserved solution composed of 0.01M PB, 0.2M ethylene glycol and 0.5M trehalose at room temperature for 2 days. The brains were embedded with OCT and cryosectioned at a temperature of -30℃∼-33℃. Brain sections were collected on cold water fish gelatin-coated slides and kept in a high humidity environment at 37℃ for 2 hours, and followed rinsed thoroughly with the 0.01M PBS for 10 minutes. Finally, brain sections were cover slipped with an on-slide clearing mounting solution containing 55% antipyrine, 0.1% DAPI (D9542, Sigma) and 1%DABCO (A51654, OKA) dissolved in the ddH2O. Before imaging, slides were kept in the dark at room temperature.

#### Parcellation and 3D reconstruction of mouse auditory thalamic nuclei

To achieve accurate connection maps of auditory corticothalamic circuits, the first thing to do is to localize and get the boundaries of brain regions composing the auditory cortex and thalamus. The Allen Mouse Brain Common Coordinate Framework Atlas (CCF)^54^ provided the most precise localization and laminar distribution of not only the auditory cortex but also sensory and sensory-related cortical areas localization and laminar discrimination. And the auditory cortex comprises the dorsal auditory area (AUDd), posterior auditory area (AUDpo), primary auditory area (AUDp) and ventral auditory area (AUDv).

It is essential to divide the subnuclei of the auditory thalamus (AT) reasonably and identify the location of viral infection. In previous studies focusing on rats, cats, and guinea pigs, the localization and division of auditory thalamic subnuclei relied on the physiological properties of neurons and histological characteristics^110–113^. Since the size and type of neurons vary across species ^114–117^, there is a need to determine the subnuclei borders for mice. However, the current mouse brain atlas does not provide an adequate tool for the registration of anatomical data with the subregions of the auditory thalamus due to incongruities with the cytoarchitecture, histochemistry, and neural physiology properties, especially for the primary part (MGBv), even the two popular mice brain atlas, the Allen Brain Atlas ^54^ and the Mouse Brain in Stereotaxic Coordinates ^118^. Based on the characteristics of dense fluorescence expression in the MGBv of Thy1-YFP mice^49,50^ and CB protein expression patterns in higher-order subnuclei of the AT^48^ (Fig S1A), we drew boundary lines between adjacent subnuclei of the AT in the coronal brain slices at different AP positions(Fig S1B). We ensured spatial rationality and universality by creating three-dimensional models of each subnucleus and obtaining their coronal maps through serial slicing of the models (Fig S1B-C). Our parcellation of the auditory thalamus was verified by other cytochemical properties or protein expression in MGBv, i.e., PV fibers ^119^ and CYO staining ^109^ (Fig S1D-E), guaranteeing the rigor of our results and providing an accurate localization and division of the mouse auditory thalamus for future research.

#### Cell-type-specific corticothalamic circuit labeling

To visualize cellular and synaptic structures, particularly axonal terminals projected to the auditory thalamus, of different auditory corticothalamic neurons, the virus AAV2/9-hSyn-flex-synaptophysin-mRuby-T2A-mGFP was injected into the auditory cortex of Ntsr1-Cre. Specifically, the mixture of multifold diluted AAV2/2-FLEX-Flpo (1000 times dilution) and AAV2/9-hSyn-fDIO-synaptophysin-mRuby-T2A-mGFP was injected into the auditory cortex of Ntsr1-Cre mice, respectively to label the proper density of neurons and avoid too dense fibers which were not conducive to morphological observation.

To achieve cell-type specificity and appropriate quantity of fluorescent labeling of auditory cortical pyramidal tract neurons (PT neurons), AAV2/9-mscRE4-Flpo, AAV2/9-fDIO-iCre with 2000-5000 dilution and AAV2/9-hSyn-flex-synaptophysin-mRuby-T2A-mGFP were injected together into the auditory cortex of wild-type mice. This virus tracing strategy allowed us to successfully obtain the projection distribution and morphological properties of axonal terminals in the auditory thalamus which were derived predominantly from auditory PT neurons.

#### Imaging and fluorescent signal recognition

In the study, different imaging techniques were combined to visualize brain tissue at varying levels of detail. For brain-wide cell counting and neuron distribution analysis, the slides were imaged with a 10x objective using Axio Scan.Z1 (Zeiss Axio Scan, German). Meanwhile, brain slices were imaged with a 20x objective using the spinning disk confocal microscopy (OXFORD Instruments ANDOR, Britain) to visualize detailed microscale structure and immunofluorescence staining. 20X objective and 5μm z-step were used for determining the colocalization of the cells. A 40X water-immersed objective and 0.5μm z-step were used for the observation and reconstruction of the precise structures of axonal fibers and terminals.

The neurons and axonal terminals were recognized in Imaris (Oxford Instrument) with characteristic diameters of 10μm and 1μm, respectively. Misidentified or missed fluorescent signals were manually supplemented or removed. Generally, the manual operation was performed in the situation where the fluorescent intensity of the proximal dendrites or axons was too high to recognize the neuron soma or discriminate adjacent neurons automatically. Finally, both the position of fluorescent points on the corresponding images and images whereon the fluorescent points signals were imported into the Rhinoceros 7 (https://www.rhino3d.com/) for registration and statistical analysis in the normalized three-dimensional brain space of mice ^120^.

#### Registration and 3D reconstruction

The areal and laminar partition of the auditory cortical fields was registered to the Allen Mouse Brain Atlas CCF^54^, and the localization of MGBv, MGBd, and other subnuclei of the auditory thalamus was identified accurately using our parcellation criteria mentioned above.

The connectivity between cortical areas and subnuclei of the auditory thalamus was defined by the number of retrogradely traced neurons and normalized as the counted number of neurons distributed in the area or the layer divided by the total number of neurons from all the auditory cortical areas.

Brain slice and the coordinates of spot signals recognized in the Imaris were imported to the Normalized 3D Mouse Brain Space (NMBS) which was developed in the Rhinoceros7 based on the .svg files of Allen Mice Atlas downloaded ^120^. A linear transformation operation was performed on the brain slices and whereon point signals to complete the registration between neural signal and their corresponding brain regions. Then, the coordinate distribution of point signals for neurons or axon terminals in the mouse brain was then obtained and documented.

The 3D coordinates ((x,y) represented the coordinates in the coronal slice, and z represented the AP distance.) of the fluorescent signal points for starter neurons and input cells for the RV tracing experiments in the whole brain were obtained by registering the brain slices with the corresponding CCF in Rhinoceros7. In case the collected brain slices did not match each standard referenced CCF or not all CCF had corresponding brain slices, the AP value of neuron coordinates in the mismatched areas was corrected to ensure consecutive and equidistant distribution of neurons in 3D space. Each point was randomly assigned z value within the +50μm∼-50μm interval of the original coordinate of the points to achieve an approximately continuous distribution of neurons on the z-axis.

#### Rendering of whole-brain input neurons’ distribution

After the coordinates (x,y,z) of dot signals were acquired in the NMBS through the registration operation described in the previous paragraph, a total number of 3000 neurons were stacked and rendered in Rhinoceros7. To display the brain-wide input distribution to the cortical pyramidal neurons of interest, 600 input neurons were randomly selected from each mouse and illustrated in the NMBS.

#### Distribution analysis of fluorescent dot signals in the cortices

To perform a quantitative analysis of the spot signal distribution in the cortices across different depths, a mapping from the temporal cortical areas to the chunking cortical space which had the shape of right cylinders was constructed in the NMBS. Firstly, the outermost surface of the temporal cortical areas which represented the pia surface of the cortex was fit by the NURBS surfaces. Using surface mapping methods, any spot with the location (x_0i_, y_0i_, z_0i_) distributed in the cortices was assigned to the corresponding coordinate (u_i_, v_i_, d_i_) in which both ui and vi were the coordinates on the NURBS surface and di represented the distance between the spot and the NURBS surface. In the transformed chunking cortical space, the coordinates (x_i_, y_i_, z_i_) of one spot were obtained by the following transformation: x_i_=u_i_, y_i_=v_i_, z_i_=d_i_/d_i0_(u_i_,v_i_). d_i0_(u_i_,v_i_) represented the actual thickness of the neocortex at a specific location (u_i_,v_i_) and was defined as the distance between the dot and the intersection of the normal line passing through the dot and the inner surface of the cortex. Therefore, the thickness of the cortices was normalized in the chunking cortical space. The neocortex was successfully transformed into right cylinders when cortical areal borders were mapped to the chunk cortical space based on the above transformation relationship. In order to analyze the density of neurons in the cortical areas from multiple groups, the dot sets with coordinates (X_0i_, Y_0i_, Z_0i_) in the NMBS were mapped into the corresponding dot sets with the coordinates (X_i_, Y_i_, Z_i_) in the chunking cortical space. Then the following adjusted quad kernel densitometry which was derived from the following formula was used for the estimation of dot signals density distribution^121^ in any location (x,y).

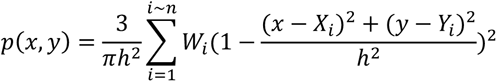

Here, Wi represented the weight of kernel and was obtained by the following expression:

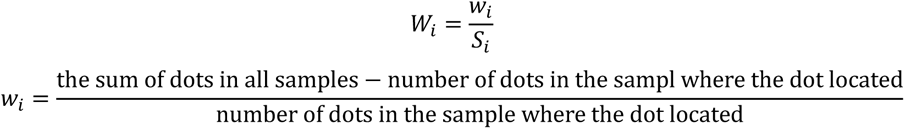

*S_i_* represented the unit area of dUdV in the NURBS.

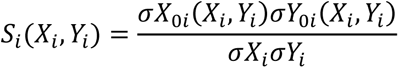

The value of h represented the bandwidth and was determined by the following rule, where Dm was the median value of the distance between the dots and their average center.

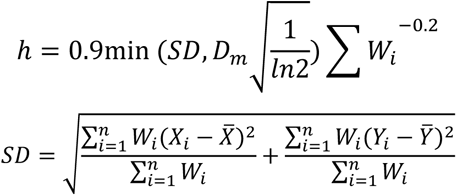

The density distribution heatmap in the chunking cortical space was obtained when the p (x, y) was assigned the corresponding color levels. Then the heatmap was projected onto the outer surface of the cortices in the NMBS using corresponding UV coordinates on the NURBS surface. Subsequently, the three-dimensional density distribution model for the fluorescent dot signals in the cortical areas was achieved after rendering in the Rhinoceros7. di in the coordinates (ui, vi, di) was used for quantifying and analyzing the depth of the dot signals within the cortex, as demonstrated in Fig 2A, D and G.

### Distribution density estimation of synaptic signals

Firstly, representative images were chosen by selecting slices with the densest projections that covered the typical coronal AP distance for brain regions of interest. Imaris Spots software was utilized to identify synaptic signals with a typical diameter of 1μm, as previously described, and the number of spot signals (n) for synaptic terminals was obtained. The density of synaptic terminals from cortical projection neurons in particular brain regions was calculated as an estimation of connection strength between cortical neurons and the brain regions according to the following equation.

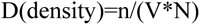

Here, the volume of the brain regions (V) was calculated as the areas of regions at the representative slices multiplied by the section thickness, which was 50μm. The number of labeled cortical neurons (N) was determined by sampling slices at intervals.

And the quantity of projections from cortical neurons to the brain regions of interest was obtained according to the following equation.

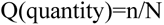

#### Quantification and analysis of terminal size acquired under light microscopy

Corticothalamic axon terminals in the AT were successfully labeled using synaptophysin-mRuby and obtained high-throughput single-synapse scale three-dimensional fluorescence imaging on large-scale brain regions using a turntable confocal microscope in junction with the original rapid and non-scaling on-slide tissue clearing method (as detailed in the methods above). The data processing path was established as depicted in Fig 4C. Firstly, deconvolution was performed according to the point spread function of the microscope system. Then, after background subtraction, the volumes of the fluorescent signals were reconstructed using the fluorescence intensity threshold. If required, adhered signals were separated by a point recognition algorithm to obtain the model of each fluorescent dot signal. Finally, the fluorescence intensity and the volume of each three-dimensional reconstructed object were extracted.

To accurately quantify the axonal terminal size under the fluorescence microscope, fluorescent microbeads with a given size (e.g., 0.5μm, 1μm, 2μm and 3μm.) that were similar in size to real synaptic terminals^122^ were utilized. This enabled us to determine the relationship between the parameters of fluorescent signals obtained by the fluorescence microscope and their real size. Microbead samples with different fluorescence intensities were imaged when injected into the agarose gel (for pre-experiments to minimize the cost of mice) or the mouse brain and their reconstructed volumes were obtained (Fig 4C1). The negative reciprocal of the mean fluorescence intensity of the microbeads and the reconstructed volume of the imaged microbeads were linearly fitted, revealing a good linear positive correlation between them (Fig 4C2-3). The relationship function between the real size of the targeted microbeads and their reconstructed volume was represented by the surface obtained through a fitting line lofted in three-dimensional space between the distribution of both the negative reciprocal of the fluorescence intensity and the reconstructed size (Fig 4C4, left). This model was used for reconstructing the volume of all the fluorescent microbeads with different sizes (0.5μm, 0.8μm, 1μm,1.5μm, 2μm and 3μm) and results showed that beads of different sizes were separated from their FI and volume and good estimations of their actual sizes were obtained. In a conclusion, our pipeline of extracting actual sizes of axonal terminals from the 3D confocal images were effective in quantifying axonal terminal size (Fig 4C5).

### QUANTIFICATION AND STATISTICAL ANALYSIS

Data are all presented as mean ± SEM. In all bar graphs, bars represent mean data and error bars represent SEM. In the trace plot, lines in darker color represent mean data, and shadows represented areas covering mean ± SEM.

Statistical tests and data analysis were performed using GraphPad Prism 8.0 (GraphPad Software, USA). All tests were two-detailed. The Shapiro-Wilk test was first applied to examine whether samples had a normal distribution. For the unpaired test of two groups, an unpaired t-test was applied in the case of normal distribution; otherwise, a non-parametric Mann-Whitney test was applied. For the paired test of two groups, the paired t-test or the Wilcoxon matched-pairs signed rank test was applied in case of normal distribution or not, respectively.

For comparison among three groups or more without pairing, ordinary one-way ANOVA and the Kruskal–Wallis test were used in the case of a normal distribution or not, respectively. For comparison among three groups or more with pairing, RM one-way ANOVA and Friedman test was used in case of normal distribution or not, respectively. In multiple comparison situations, Tukey’s and Dunn’s were chosen as post hoc tests in the situation of normal distribution and otherwise, respectively. In case of unequal SDs when comparing among three groups or more without pairing, Brown-Forsythe ANOVA and Welch’s ANOVA tests with Tamhane’s T2 multiple comparisons test as post hoc test were used. If multiple comparisons were performed between a particular dataset with all the other datasets, Dunnett’s multiple comparisons test was used in the case of a normal distribution. For comparisons with two independent variables, ordinary two-way ANOVA followed by Tukey’s multiple comparisons test as post hoc test depending on the factors to be tested. For comparisons of two groups for each row, multiple unpaired t tests with Holm-Sidak method as post hoc tests were used. Significance was set at *P*<0.05 and was indicated as \**P*<0.05, \*\**P*<0.01, \*\*\**P*<0.001, \*\*\*\**P*<0.0001, n.s.*P*>0.05. (Note: some not specifically marked). Details of each statistical test used were described in the figure legends and listed in Table S2(related figure, sample size, normality and variance situation, statistic method, statistics value, P values, and necessary post hoc multiple comparison test results).

---

**Figure S1.**
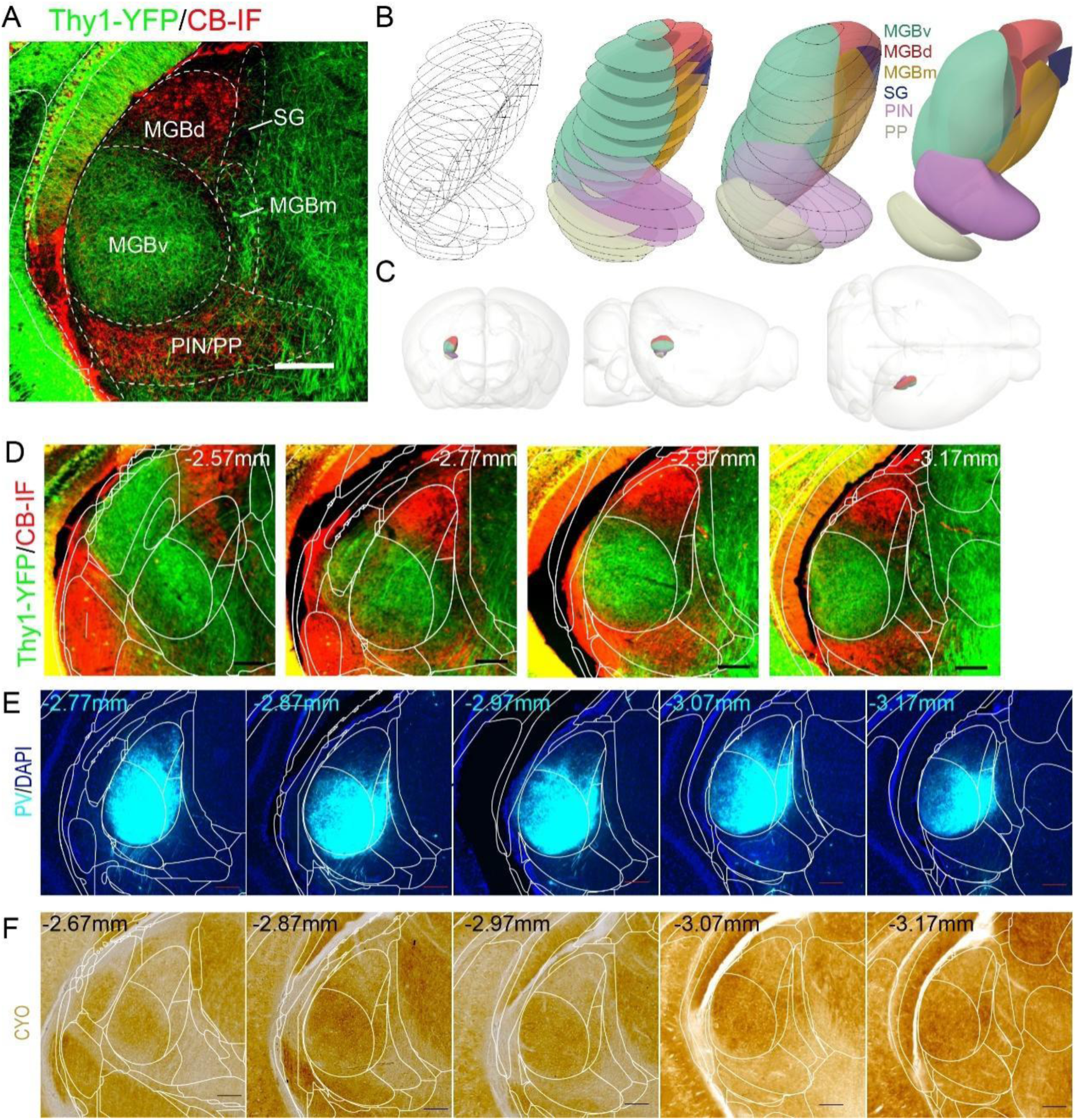
Extended data to Figure 1. Parcellation, conduction of 3D modeling and 2D atlas slicing for mouse auditory thalamus. (A) Division and annotation of auditory thalamic subnuclei based on YFP distribution and CB immunofluorescence results in the Thy1-YFP mice. Scale bar, 200μm. (B) Process of 3D modeling of auditory subnuclei. Wireframes were derived from parcellation of YFP and CB immunostaining results. (C) Three different views of the auditory thalamus in the mouse brain 3D model. (D) Borders between adjacent auditory subnuclei at different AP distances from the Bregma in the mouse coronal brain sections. Scale bars, 200μm. (E) Good fit between PV fiber distribution and the coronal atlas sectioned from auditory thalamic 3D model at the corresponding AP distances. Scale bars, 200μm. (F) CYO staining results matching the coronal atlas sectioned from auditory thalamic 3D model at the corresponding AP distances. Scale bars, 200μm.

**Figure S2.**
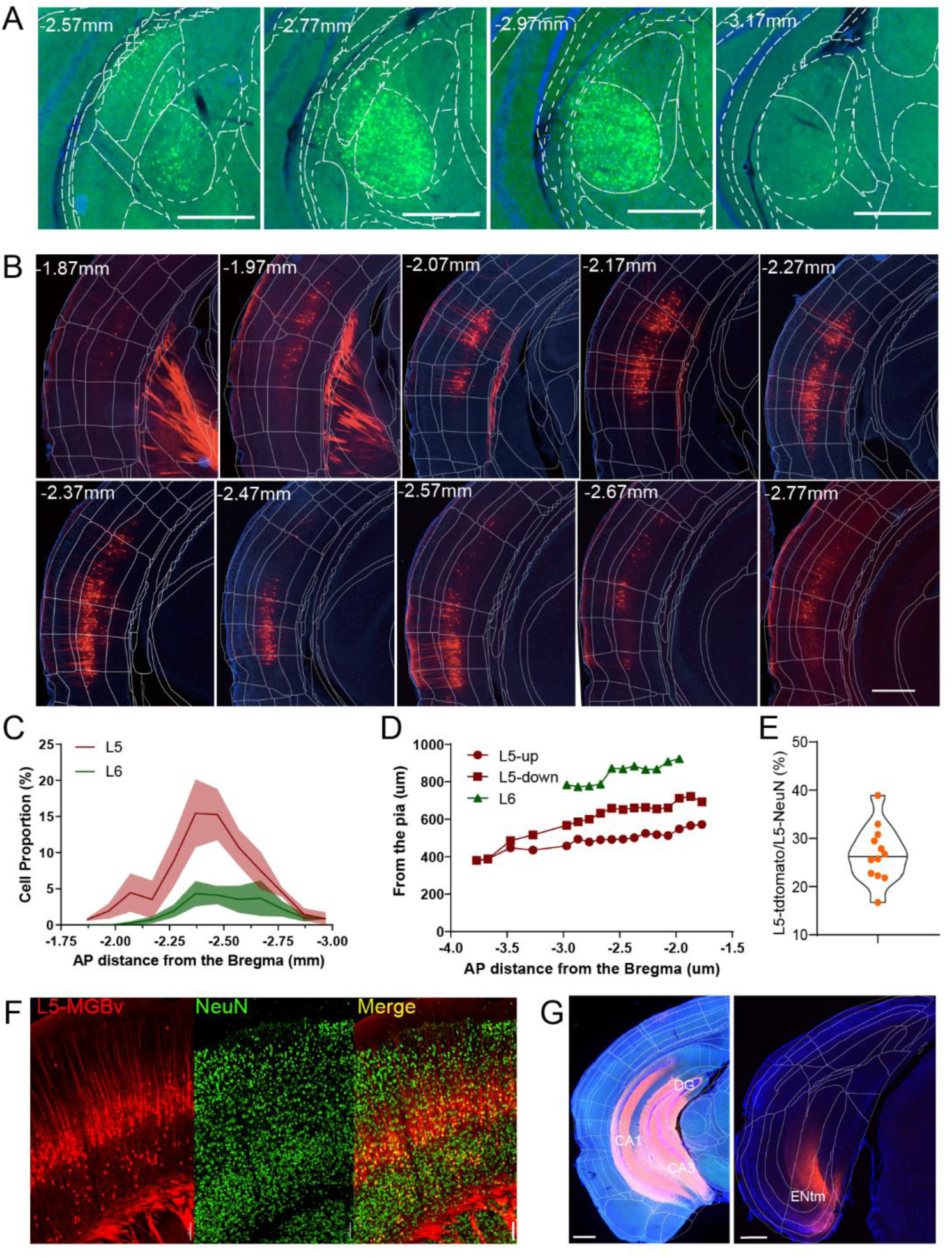
Extended data to Figure 1. Auditory L5 neurons projecting to MGBv were retrogradely labeled by rAAV2-retro in the auditory cortex. (A-B) Representative images of MGBv (A) locally transfected by rAAV2-retro and the auditory cortical areas expressing tdTomato (B) obtained from coronal brain slices of the same mouse. Scale bars, 500μm. (C) Distribution proportions of L5 MGBv-projecting and L6 MGBv-projecting neurons in the auditory cortex along the anterior-posterior axis. Data were from 5 mice and are presented as mean (dark lines)±SEM(shadow). (D) Depth distribution of retrogradely L5 and L6 MGBv-projecting neurons at the different AP distances from the Bregma. L5-up neurons indicate the shallowest and deepest locations of L5 MGBv-projecting neurons, respectively. (E) Proportions of L5 MGBv-projecting neurons labeled by rAAV2-retro in all L5 neurons labeled by NeuN. Data were from 10 brain slices of 2 mice. (F) Representative confocal image of L5 MGBv-projecting neurons labeled by rAAV2-retro (Red) and NeuN+ cells (Green) in the auditory cortex. Scale bars, 100μm. (G) Distribution of labeled cortical neurons (right panel) after AAV2-retro injection into the hippocampus (left panel) along the injection trace. Scale bars, 500μm.

**Figure S3.**
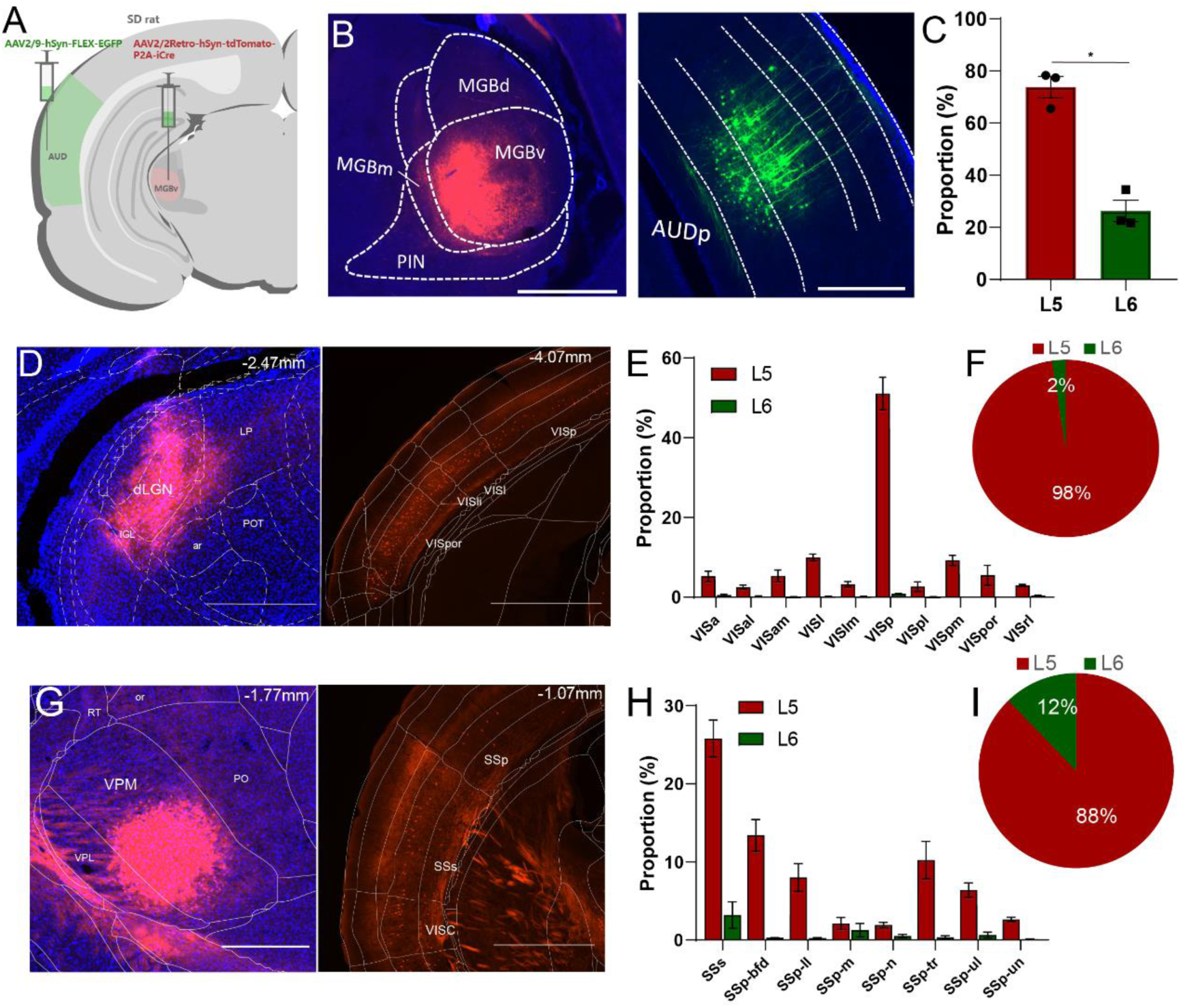
Extended data to Figure 1. L5 neurons project to the first-order sensory thalamus in the auditory system of rats and visual and somatosensory systems of mice. (A) Schematic of retrograde tracing for the MGBv-projecting cortical neurons in rats by rAAV2-retro. (B) Representative images of the location of rAAV2-retro represented by mCherry concentrated in the rat MGBv (left) and corresponding retrogradely labeled MGBv-projecting cortical neurons indicated by GFP in the rat auditory cortex (right). Scale bars, 1000μm. (C) Proportions of L5→MGBv neurons and L6→MGBv neurons retrogradely labeled by rAAV2-retro in auditory cortex of rats (n=3). Statistic method was paired t test. P value, * *P*<0.05. (D) Local injection of rAAV2-retro-Cre was performed in the dLGN (left) and retrograde labeling of dLGN-projecting neurons by rAAV2-retro-Cre in the visual cortex of mice (right). Scale bars in left panel,500μm. Scale bars in right panel,1000μm. (E) The proportions of dLGN-projecting neurons across different visual cortical areas. Data are presented as mean±SEM and were obtained from 4 mice. (F) Pie chart showing the proportions of L5 and L6 in the dLGN-projecting neurons. (G) Local injections of rAAV2-retro-Cre were performed in the VPM (left) and retrograde labeling of VPM-projecting neurons in the somatosensory cortex of mice (right). Scale bars in left panel, 500μm. Scale bars in right panel,1000μm. (H) The proportions of VPM-projecting neurons across different visual cortical areas. Data are presented as mean±SEM and were from 5 mice. (I) Pie chart showing the proportions of L5 and L6 in the VPM-projecting neurons.

**Figure S4.**
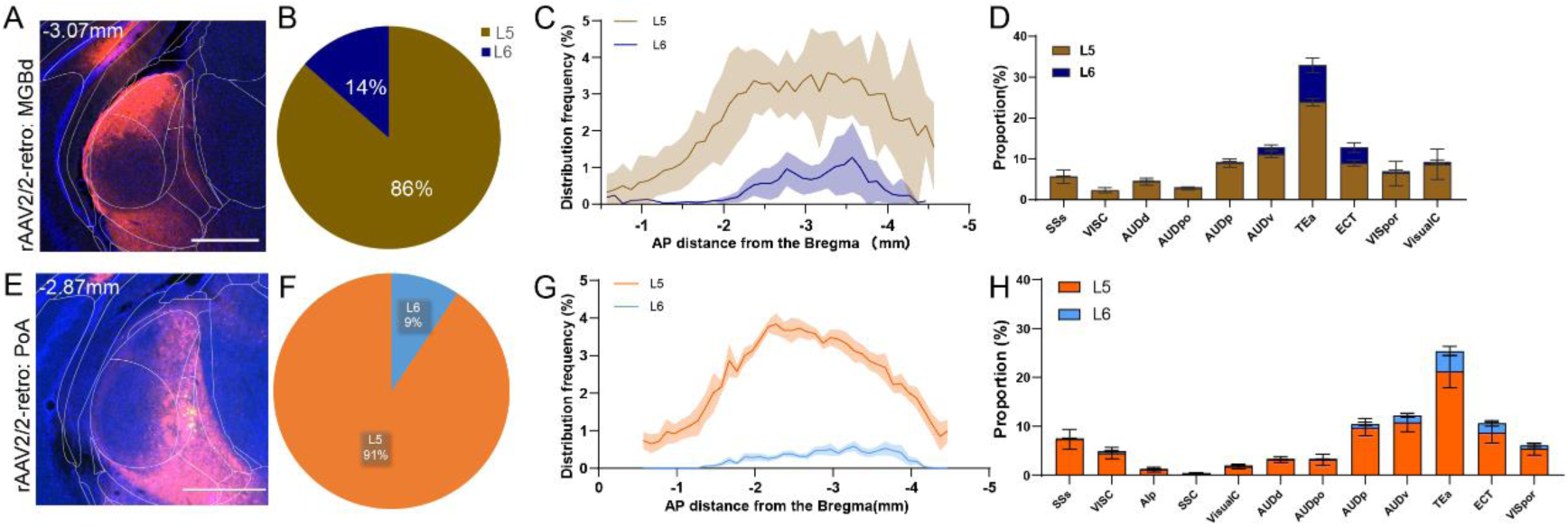
Extended data to Figure 2. rAAV2-retro preferentially retrogradely labeled L5 neurons projecting to the non-primary and polymodal association modules of the AT. (A&E) Representative images of coronal brain slices showing rAAV2-retro was locally injected into the MGBd (A) and PoA (E) of Ai14 micec, respectively. Scale bars, 500μm. (B-D) Proportions of L5→MGBd and L6→MGBd neurons retrogradely labeled by rAAV2-retro in total (B), at different AP distances from the Bregma (C) and in different cortical regions (D). Data were from 6 mice and are presented as mean±SEM. (F-H) Proportions of L5→PoA and L6→PoA neurons retrogradely labeled by rAAV2-retro in total (F), at different AP distances from the Bregma (G) and in different cortical regions (H). Data were from 7 mice and are presented as mean±SEM.

**Figure S5.**
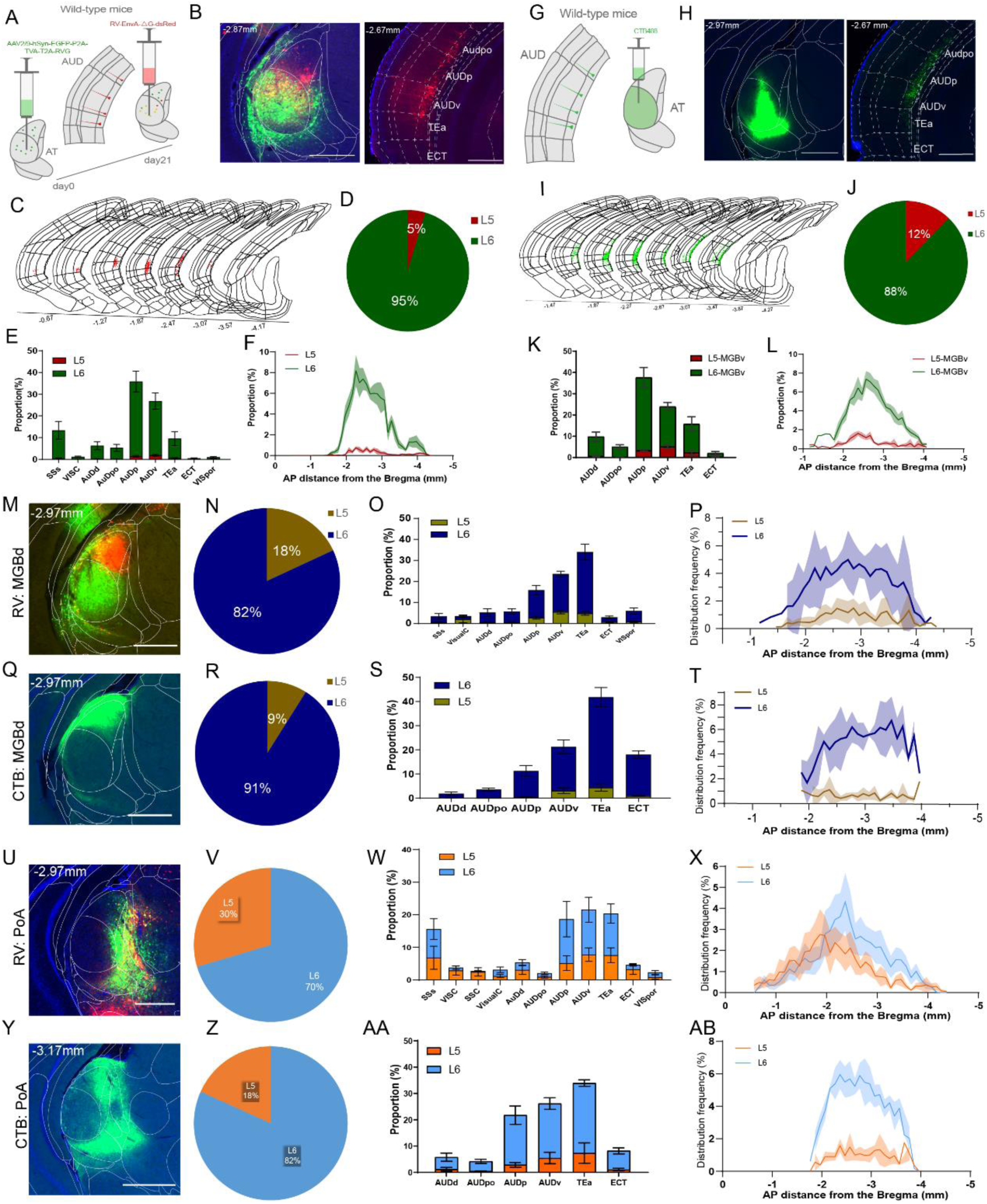
Extended data to Figure 2. RV and CTB preferentially retrogradely labeled L6 neurons projecting to different modules of the auditory thalamus. (A) Schematic representation of RV with the adeno-associated viruses as helper injected in the auditory thalamus of mice. (B) Example brain slice images for retrograde tracing of MGBv using RV strategy. Left panel shows start neurons locally distributed in MGBv. Right panel shows retrograde labeling of neurons in auditory cortical areas. Green represents TVA-GFP. Red represents RV-dsRed. Orange represents start neurons. Blue represents DAPI. Scale bars, 500μm. (C) Typical distribution of RV-labeled MGBv-projecting neurons in the coronal view of auditory cortices. (D) Proportions of L5 and L6 neurons retrogradely labeled when RV starter neurons were locally concentrated in MGBv. Data were from 6 mice, same as E-F. (E-F) Proportions of RV-labeled L5 and L6 MGBv-projecting neurons distributed in different cortical regions (E) and at the different AP distances from the Bregma (F). Green line represents L6, while red line represents L5. Data are presented as mean±SEM. (G) Schematic representation of CTB injection in the auditory thalamus of mice. (H) Example brain slice images of MGBv retrograde tracing using CTB. Left panel shows CTB injection concentrated in MGBv. Right panel shows retrograde labeling of neurons in auditory cortical areas. Green represents CTB and blue represents DAPI. Scale bars, 500μm. (I) Typical distribution of CTB-labeled MGBv-projecting neurons in the coronal view of auditory cortices. (J) Proportions of L5 and L6 neurons retrogradely labeled when CTB was locally concentrated in MGBv. Data were from 6 mice, same as K-L. (K-L) Proportions of CTB-labeled L5 and L6 MGBv-projecting neurons distributed in different cortical regions (K), and at the different AP distances from the Bregma (L). Green line represents L6, while red line represents L5. Data are presented as mean±SEM. (M, Q, U&Y) Representative images of coronal brain slices showing MGBd and PoA locally transfected by RV (M for MGBd and U for PoA) and CTB (Q for MGBd and Y for PoA) respectively. Scale bars, 500μm. (N-P) Proportions of L5→MGBd and L6→MGBd neurons retrogradely labeled by RV in total number (N), in different cortical regions (O) and at different AP distances from the Bregma (P). Data were from 6 mice and are presented as mean±SEM in O-P. (R-T) Proportions of L5→MGBd and L6→MGBd neurons retrogradely labeled by CTB in total number (R), in different cortical regions (S) and at different AP distances from the Bregma (T). Data were from 3 mice and are presented as mean±SEM in S-T. (V-X) Proportions of L5→PoA and L6→PoA neurons retrogradely labeled by RV in total number (V), in different cortical regions (W) and at different AP distances from the Bregma (X). Data were from 7 mice and are presented as mean±SEM in W-X. (Z-AB) Proportions of L5→MGBd and L6→MGBd neurons retrogradely labeled by CTB in total number (Z), in different cortical regions (AA) and at different AP distances from the Bregma (AB). Data were from 3 mice and are presented as mean±SEM in AA-AB.

**Figure S6.**
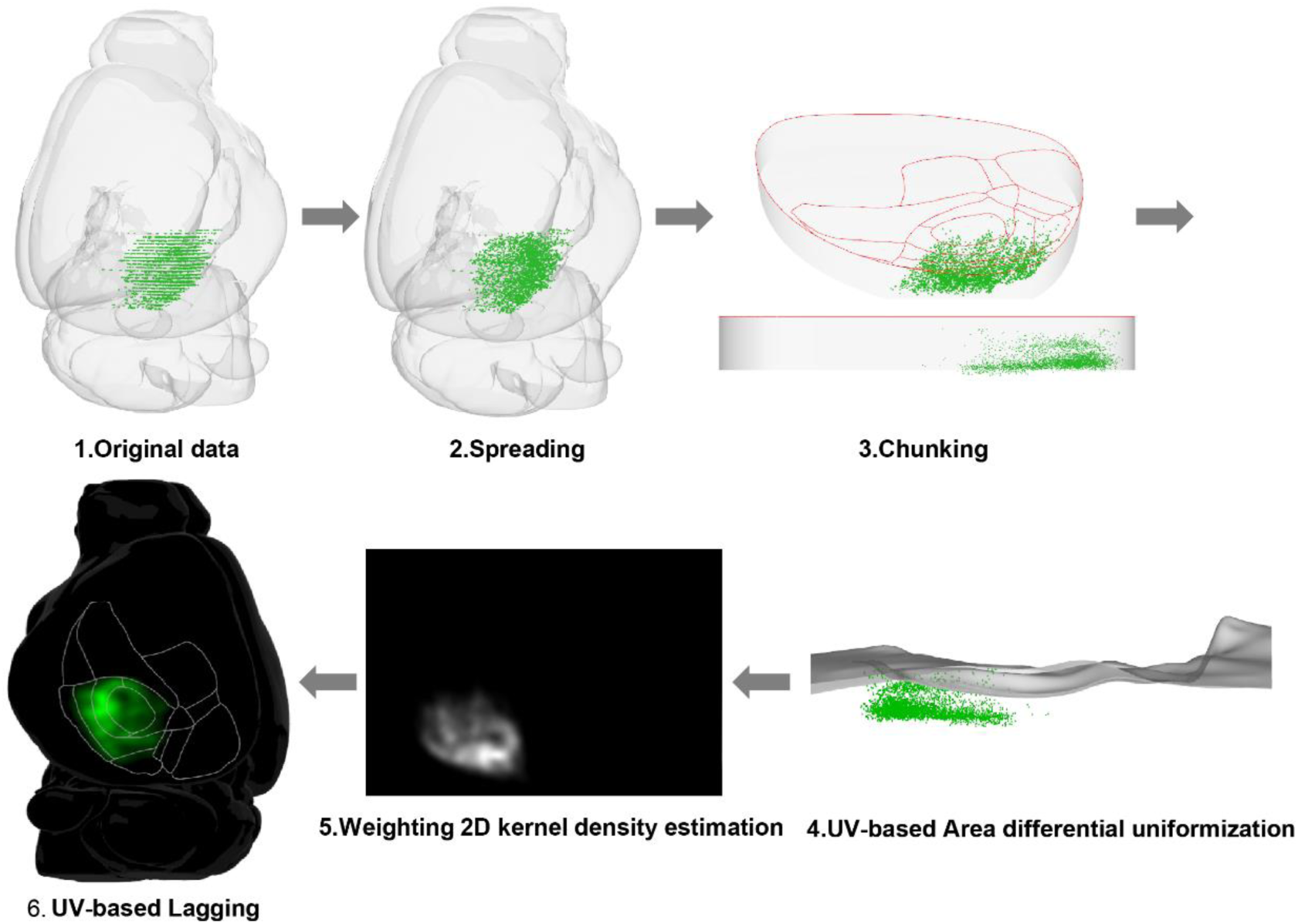
Extended data to Figure 2. Density distribution estimation of AT- projecting cortical neurons by KDE analysis.

**Figure S7.**
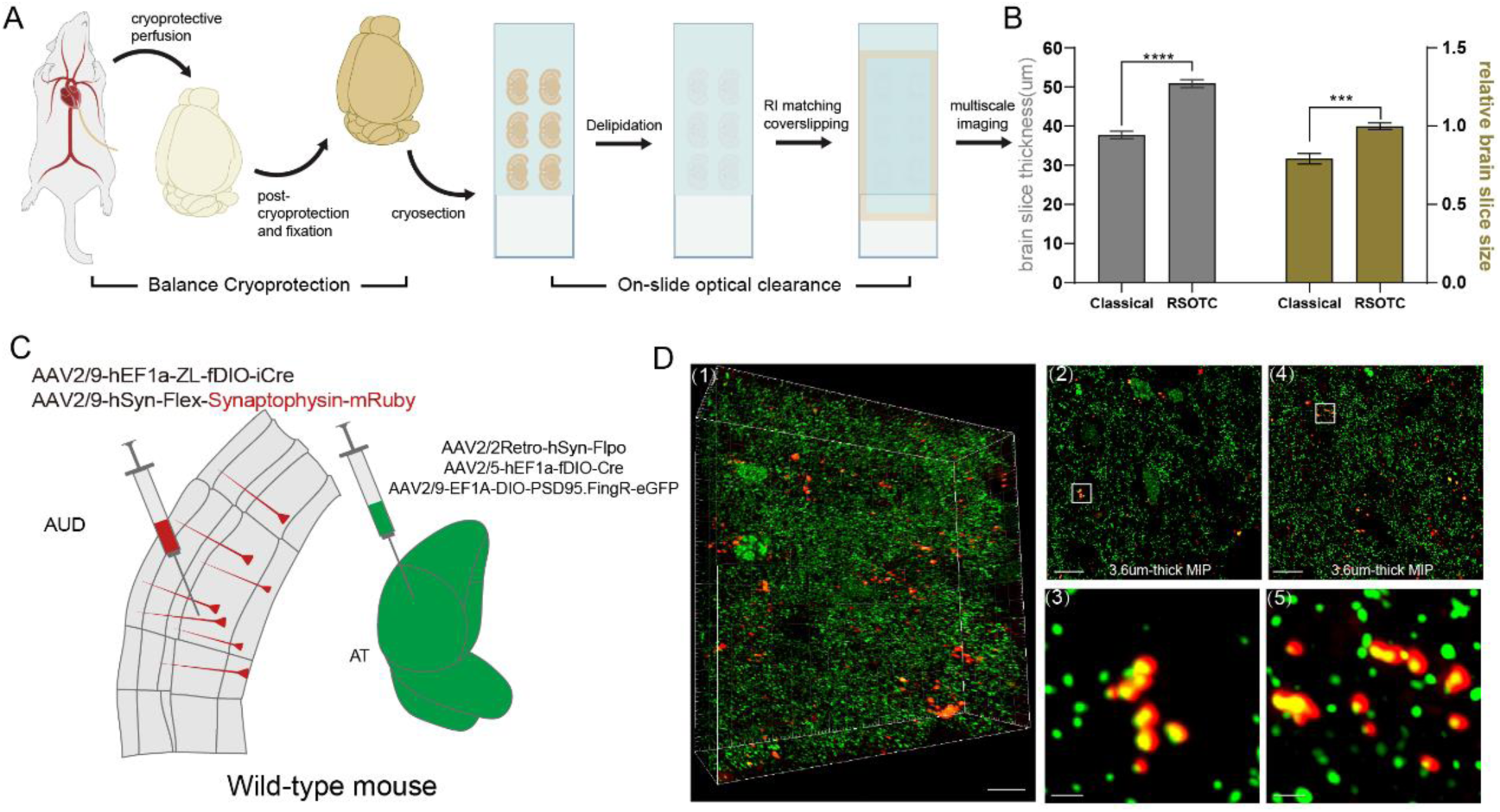
Extended data to Figure 3. Rapid on-slice clearing for microscale synaptic connection between synaptophysin and PSD95. (A) Overview of the brain sectioning and optical clearing before imaging. (B) Comparison of change in tissue size after processing between traditional clearing and our developed RSOTC. Data were from 7 samples. Statistical methods were both unpaired t-test. P value, *** *P*<0.001, **** *P*<0.0001. (C) The schematic of viral injection strategy to label auditory corticothalamic terminals and PSD95 of auditory thalamic neurons. (D) Confocal images showing the synapse pairs between the presynaptic terminals of retrogradely labeled auditory corticothalamic neurons (red) and PSD95 of neurons in the auditory thalamus (green). D1, the selected voxels containing synaptic pairs of corticothalamic circuits. Images D2-D3 and D4-D5 represent synaptic pairs from two different sites in the D1. Scale bars in (D1-D2) and (D4), 10μm. Scar bars in (D3) and (D5), 1μm.

**Figure S8.**
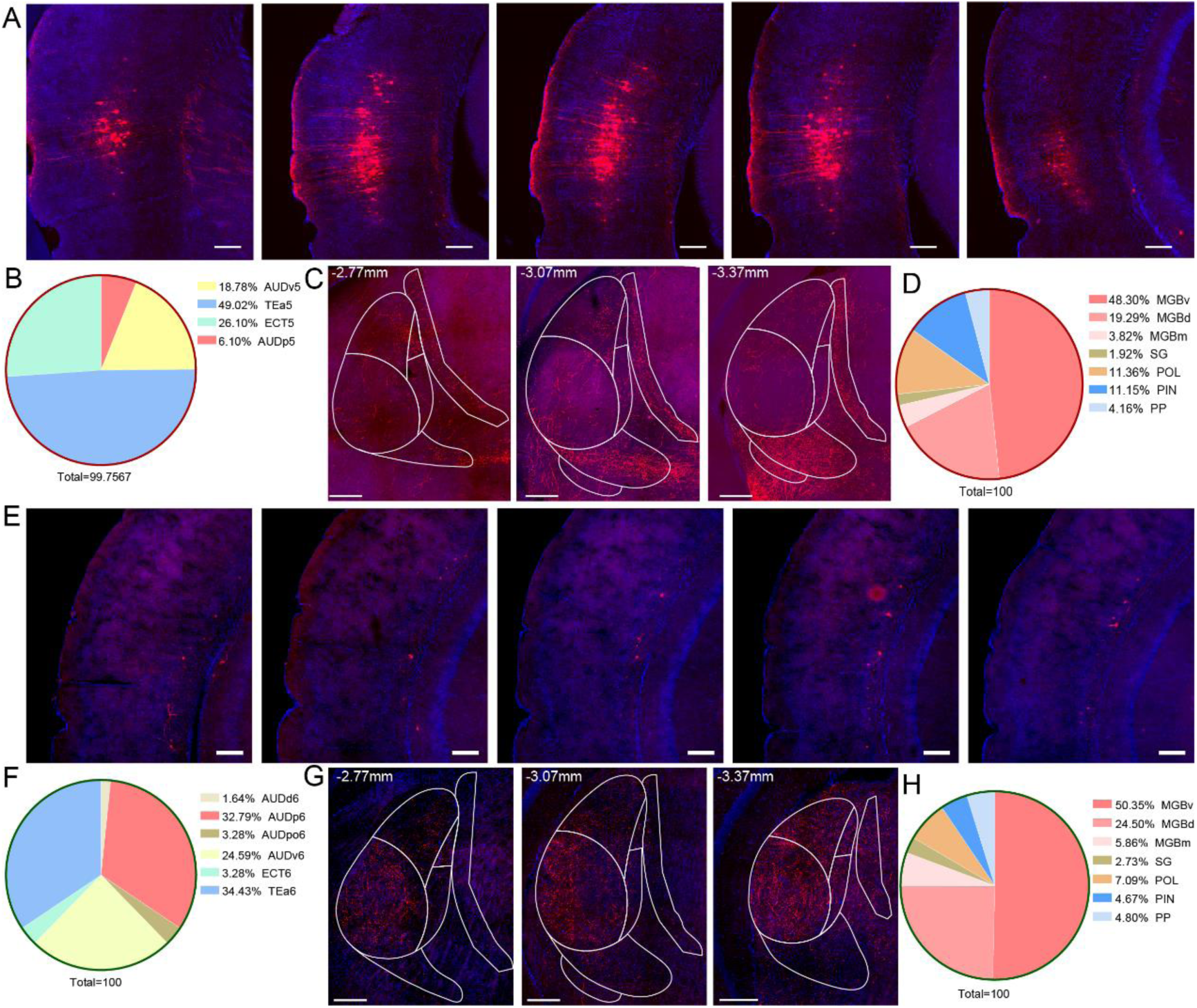
Extended data to Figure 3. Distribution of axon terminal exclusively from rL5→AT and rL6→AT neurons in the auditory thalamus. (A-B) Confocal images showing exclusive retrogradely sparse labeling of L5 AT-projecting neurons in the auditory cortices and proportions of these neurons located in different auditory-related cortical areas. Scale bars, 200μm. (C-D) Confocal images displaying axon terminals derived from L5 AT-projecting neurons (C) and distribution proportions of these terminals in distinct subnuclei of the auditory thalamus (D). Scale bars, 200μm. (E-F) Confocal images showing exclusive retrogradely sparse labeling of L6 AT-projecting neurons in the auditory cortices and proportions of these neurons located in different auditory-related cortical areas. Scale bars, 200μm. (G-H) Confocal images displaying axon terminals derived from L6 AT-projecting neurons and distribution proportions of these terminals in distinct subnuclei of the auditory thalamus. Scale bars, 200μm.

**Figure S9.**
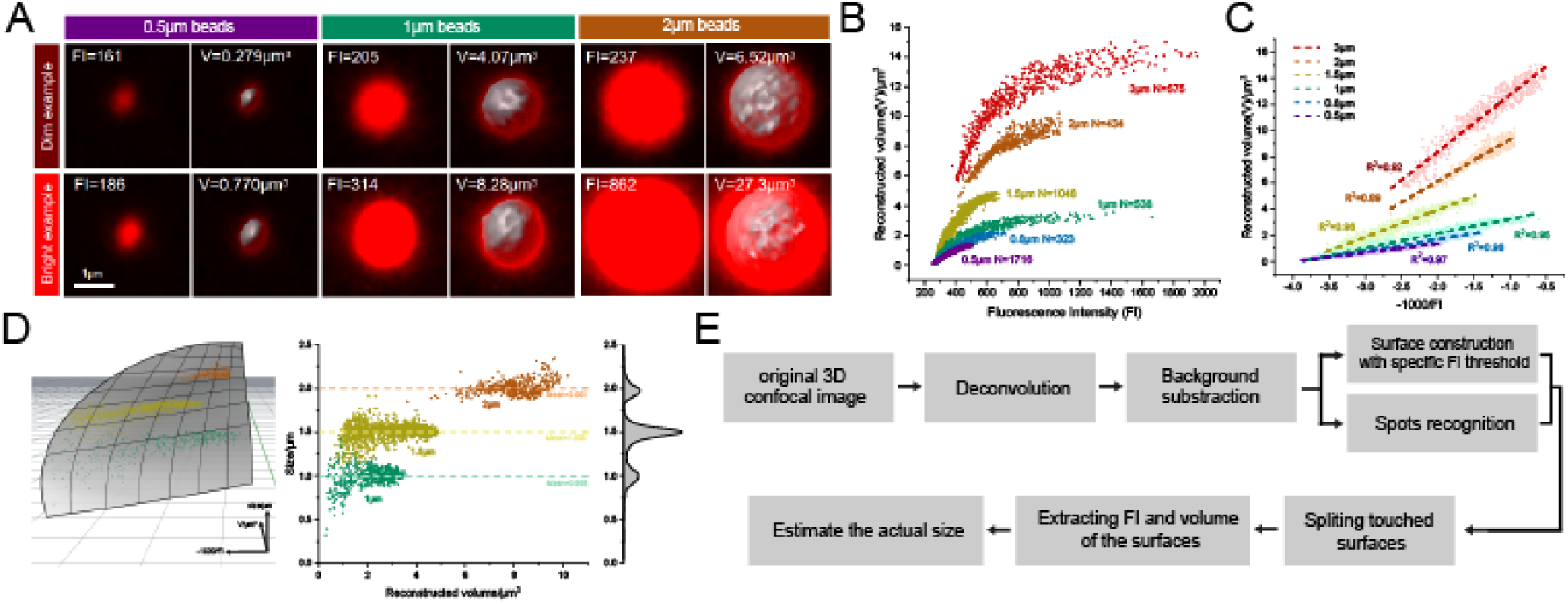
Extended data to Figure 4. Stereological method for quantitative analysis of fluorescent signals at axonal terminals. (A) Reconstruction of fluorescent beads with known sizes using closed surfaces. The volume (V) of these surfaces is relevant to their mean fluorescent intensity (FI). Although beads of the same size, brighter ones are reconstructed with surfaces of larger volume. (B-C) Linear correlation between the reciprocal of FI and the reconstructed volume, regardless of bead size. Beads of different sizes fall into distinguishable linear functions. (D) Left, a surface is constructed to fit the relationship among mean fluorescent intensity, reconstructed volume, and actual size. The beads of 1μm, 1.5μm, and 2μm are projected onto this surface. Right, the fitting method separates beads of different sizes based on their FI and volume and accurately estimates their actual sizes. (E) Pipeline of extracting actual sizes of axonal terminals from 3D confocal images. The fitting surface, which refers to beads of known sizes, is used to estimate the size of terminals in the last step.

**Figure S10.**
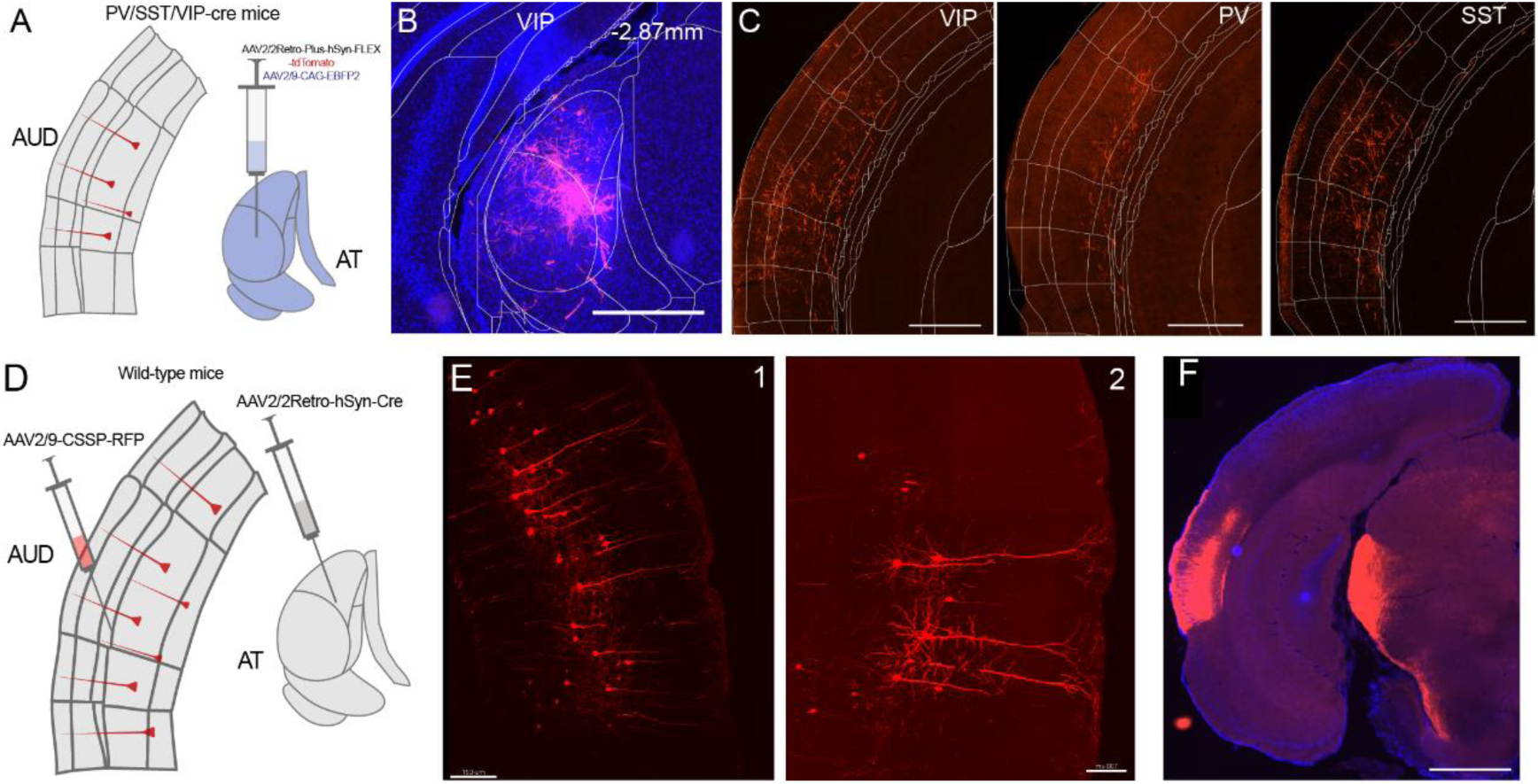
Extended data to Figure 5. Cell type validation of rL5→AT and rL6→AT neurons. (A) Schematic diagram illustrating the specific labeling of AT-projecting neurons in PV-Cre, SST-Cre, and VIP-Cre mice. (B-C) Representative retrograde labeling results showing that after rAAV2-retro-Flex-tdTomato injection into the AT (B), there were not any labeled VIP+ (C-left panel), PV+ (C-middle panel), or SST+ (C-right panel) neurons in auditory cortices. Scale bars in B-C, 500μm. (D) Schematic illustrating the sparse labeling of AT-projecting neurons in the auditory cortex for single-cell morphological analysis. (E) Confocal images showing the sparse and highlighted labeling of rL5→AT neurons in the auditory cortex. Scale bar in left panel, 150μm. Scale bar in right panel, 100μm. (F) Representative wide-field image showing pyramidal tract from rL5→AT neurons. Scale bar, 1000μm.

**Figure S11.**
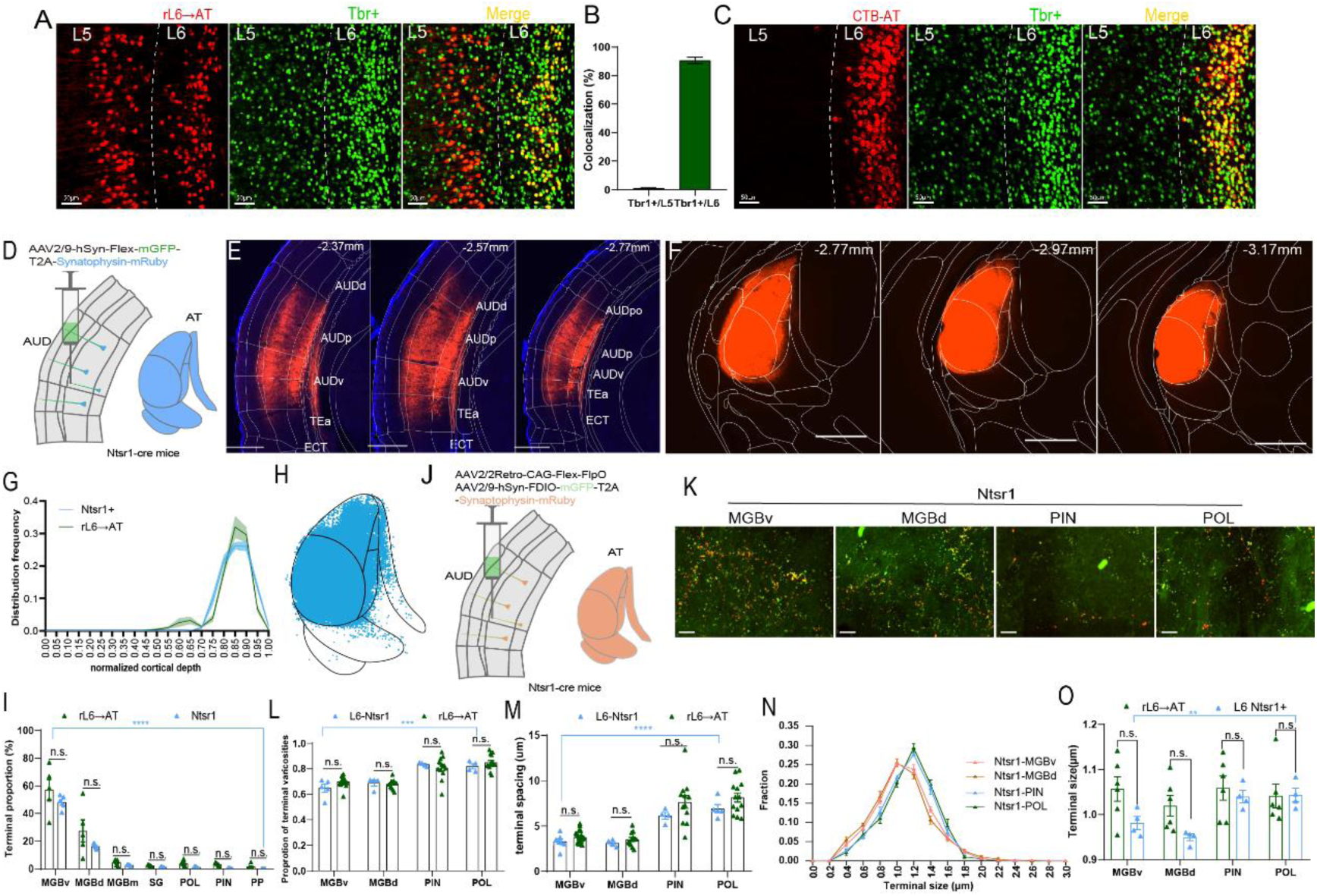
Extended data to Figure 5. rL6→AT neurons exhibited similar corticothalamic projections with L6 CT neurons. (A) Immunofluorescent staining of Tbr1 on the auditory cortex showing retrogradely labeled L5 and L6 AT- projecting neurons by rAAV2-retro. Red, AT-projecting neurons expressing tdTomato. Green, Tbr1 positive neurons. Orange, colocalizations of AT-projecting neurons with Tbr1 positive neurons. Scale bars, 50μm. (B) Proportions of rAAV2-retro-labeled AT-projecting neurons expressing Tbr1. Data were mean±SEM and collected from 7 slices of 2 mice. (C) Immunofluorescent staining of Tbr1 on the auditory cortex showing L6 AT-projecting neurons retrogradely labeled by CTB. Red, AT-projecting neurons expressing Alexa555. Green, Tbr1 positive neurons. Orange, colocalizations of L6 AT-projecting neurons with Tbr1 positive neurons. Scale bars, 50μm. (D) Schematic of fluorescent-labeled auditory cortical Ntsr1+ neurons and their axon terminals. (E-F) Representative confocal images showing Ntsr1+ neurons in the auditory cortex (E) and axon terminals from them in the auditory thalamus (F). Red fluorescence, synaptophysin-mRuby. Scale bars, 500μm. (G) Illustration of distribution frequency of Ntsr1+ neurons and rL6→AT neurons along the normalized depth of auditory cortices. (H) Three thousand points which were distributed in the auditory thalamus representing axon terminals of Ntsr1+ neurons in the auditory cortex from weighted samples. (I) Comparisons of terminal proportions across the AT subdivisions between projections of auditory Ntsr1+ neurons and those of rL6→AT neurons. Data were both from 5 mice and are presented as mean±SEM. (J) Schematic illustrating strategy of viral injection to specifically label auditory Ntsr1+ axon terminals with relative sparse degree. (K) Representative confocal images displaying axons (green) and terminals (overlap of green and red) distributed in MGBv (I1), MGBd (I2), PIN (I3), and POL (I4) from Ntsr1+ neurons in the auditory cortex. Scale bars, 10μm. (L-M) Comparisons of proportions of varicosity-type terminals (L) and terminal spacing (M) from Ntsr1+ neurons and those from rL6→AT neurons to the MGBv, MGBd, PIN and POL. Data are presented as mean±SEM. Data were from 4 mice for Ntsr1+ projections and 6 mice for rL6→AT projections same as Fig 4. Imaging field numbers for Ntsr1: 7 in MGBv, 5 in MGBd, 5 in PIN, and 6 in POL. Field numbers for Ntsr1, 5 in MGBv, 4 in MGBd, PIN, and POL. (N) Frequency distribution of axon terminal sizes from auditory L6 Ntsr1+neurons to MGB including MGBv and MGBd, as well as PIN and POL. Sample sizes: MGBv (76468 terminals), MGBd (48527), PIN (3113), and POL (2401) from 4 mice. (O) Comparisons of terminal sizes between axon terminals from L6 Ntsr1+ neurons and those from rL6→AT neurons to subnuclei of AT including MGBv, MGBd, PIN and POL. Multiple unpaired t tests with correction for using the Holm-Sidak method were used for comparisons in the same AT subdivisions between Ntsr1+ and rL6→AT. For comparisons within Ntsr1+ or rL6→AT across AT subdivisions, RM one-way ANOVA with Dunnett’s multiple comparisons test, Kruskal-Wallis test with Dunn’s multiple comparisons test, ordinary one-way ANOVA with Tukey’s multiple comparisons test, and RM one-way ANOVA with Tukey’s multiple comparisons test were performed in I, L, M and O, respectively. P values, n.s. *P*>0.05, \*\**P*<0.01, \*\*\*\**P*<0.0001. More details in Table S2.

**Figure S12.**
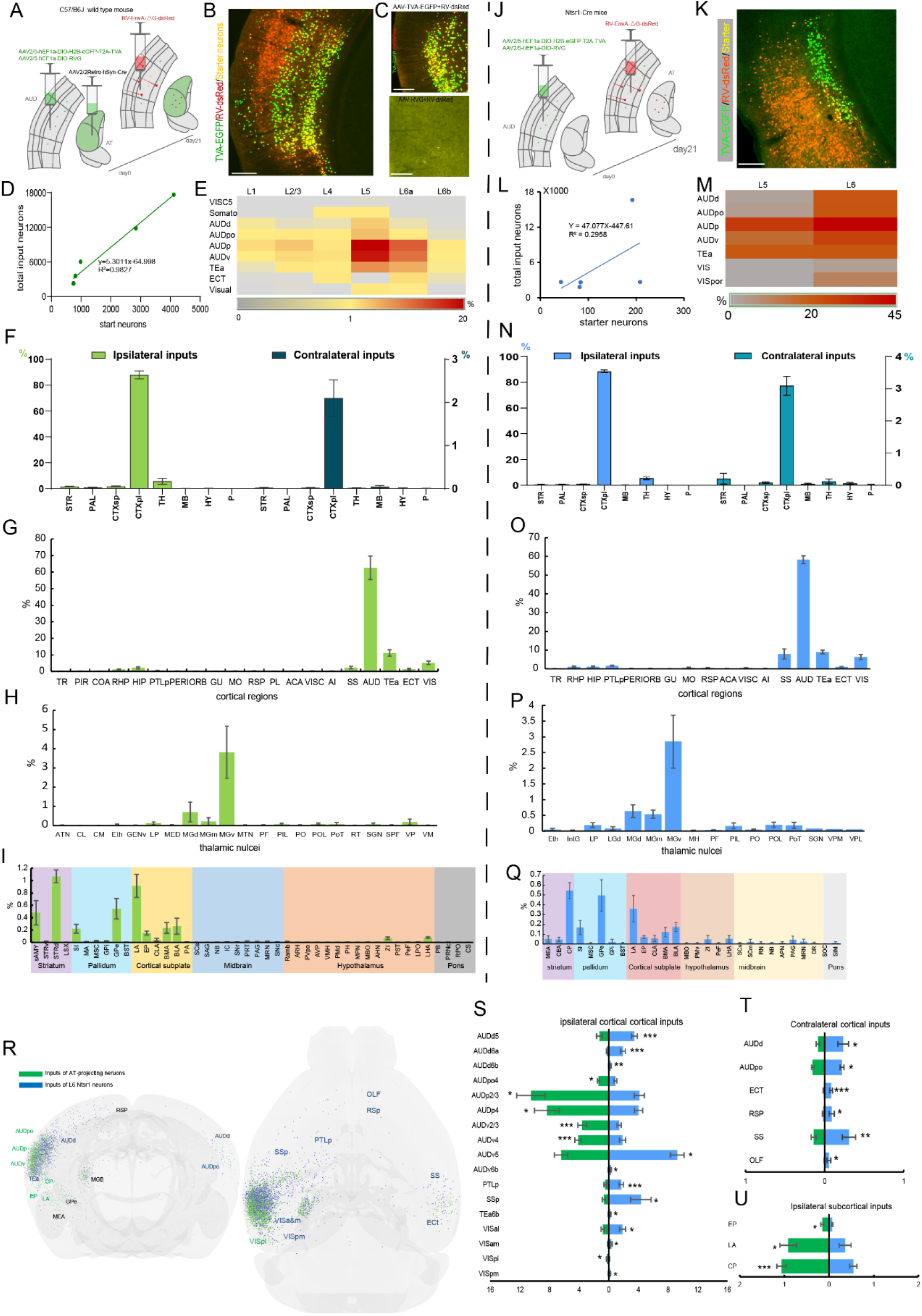
Whole-brain inputs of rAAV2-retro labeled PT neurons and Ntsr1 CT neurons. (A) Schematic showing tracing of input neurons of rAAV2-retro-labeled PT AT-projecting neurons in the auditory cortex. (B) Representative brain slice displaying start neurons and input neurons of auditory rAAV2-retro-labeled PT AT- projecting neurons. Scale bar, 200μm. (C) Brain slice obtained from the control experiments where colocalization of TVA-eGFP and RV-dsRed neurons without RVG injection (top panel) and none of neurons were labeled in the cortex when without TVA-eGFP injection (bottom panel). Scale bars, 200μm. (D) Correlation analysis between the number of starter neurons and the total number of input neurons. Data were from 5 mice. Text in the corresponding colors indicates the linear fit result and quality. (E) Heatmap showing the distribution of starter neurons of rAAV2-retro-labeled PT AT-projecting neurons in the auditory cortex. Data are presented as mean. (F-I) Distribution percentages of input neurons across different brain regions in the ipsilateral and contralateral hemispheres (F), ipsilateral cortical regions (G), thalamic nuclei (H), and nuclei in other six different subcortical regions (I). Data are presented as mean±SEM. (J) Schematic illustrating the viral tracing strategy for transsynaptic retrogradely tracing inputs to auditory L6 Ntsr1+ neurons using RV. (K) Representative brain slice showing the start neurons and input neurons of auditory L6 Ntsr1+ neurons. Scale bar, 200μm. (L) Correlation analysis between the number of starter neurons and the total number of input neurons of auditory L6 Ntsr1+ neurons. Data were from 5 mice. Text in the corresponding colors indicates the linear fit result and quality. (M) Heatmap quantifying the distribution of starter neurons of L6 Ntsr1+ neurons in the auditory cortex. Data are presented as mean. (N-Q) Distribution percentages of input neurons of auditory L6 Ntsr1+ neurons across different brain regions in the ipsilateral and contralateral hemispheres (N), ipsilateral cortical regions (O), thalamic nuclei (P), nuclei in other six different subcortical regions (Q). Data are presented as mean±SEM. (R) Input neuron distributions of PT AT-projecting neurons retrogradely labeled by rAAV2-retro and L6 Ntsr1+ neurons in the NMBS, respectively. Green dots represent input neurons of AT-projecting neurons, and blue dots represent input neurons of Ntsr1+ neurons. Green text indicates that the proportion of AT-projecting neurons receiving input from this brain area is higher than that of L6 Ntsr1+ neurons. Blue text indicates that proportion of L6 Ntsr1+ neurons receiving input from this brain area is higher than that of AT-projecting neurons. And black text indicates that there is no difference in the proportion of inputs neurons between AT-projecting neurons and L6 Ntsr1+ neurons. (S-U) Ipsilateral cortical regions (S), contralateral cortical regions (T), and ipsilateral subcortical nuclei (U) with a significant difference between input neurons of auditory PT AT-projecting neurons and L6 Ntsr1+ neurons. Statistical methods used were unpaired t test. P value: * *P*<0.05, ** *P*<0.01, *** *P*<0.001. More details in Table S2.

**Table S1.**
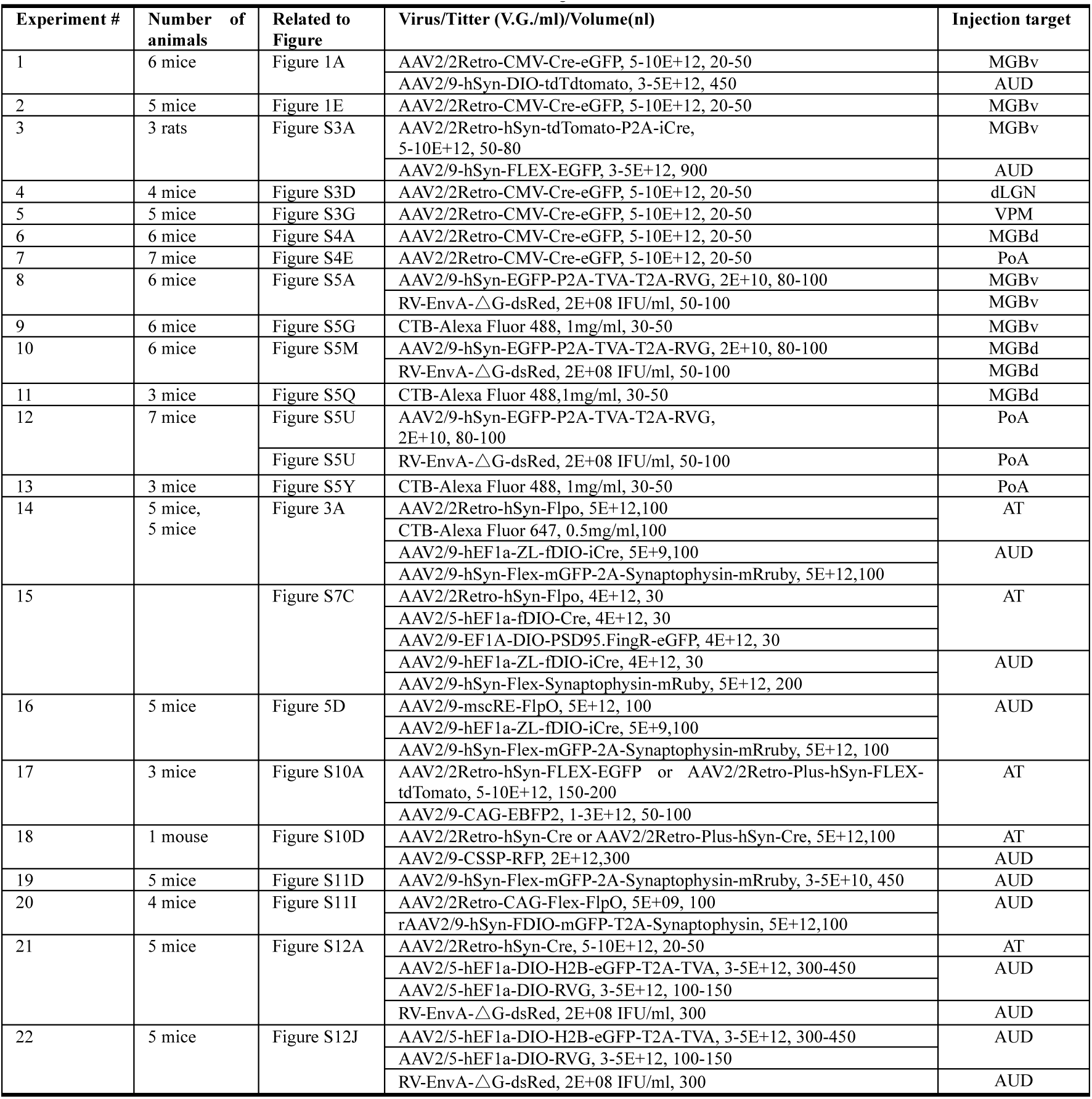
Detailed information on virus injection.

**Table S2.**
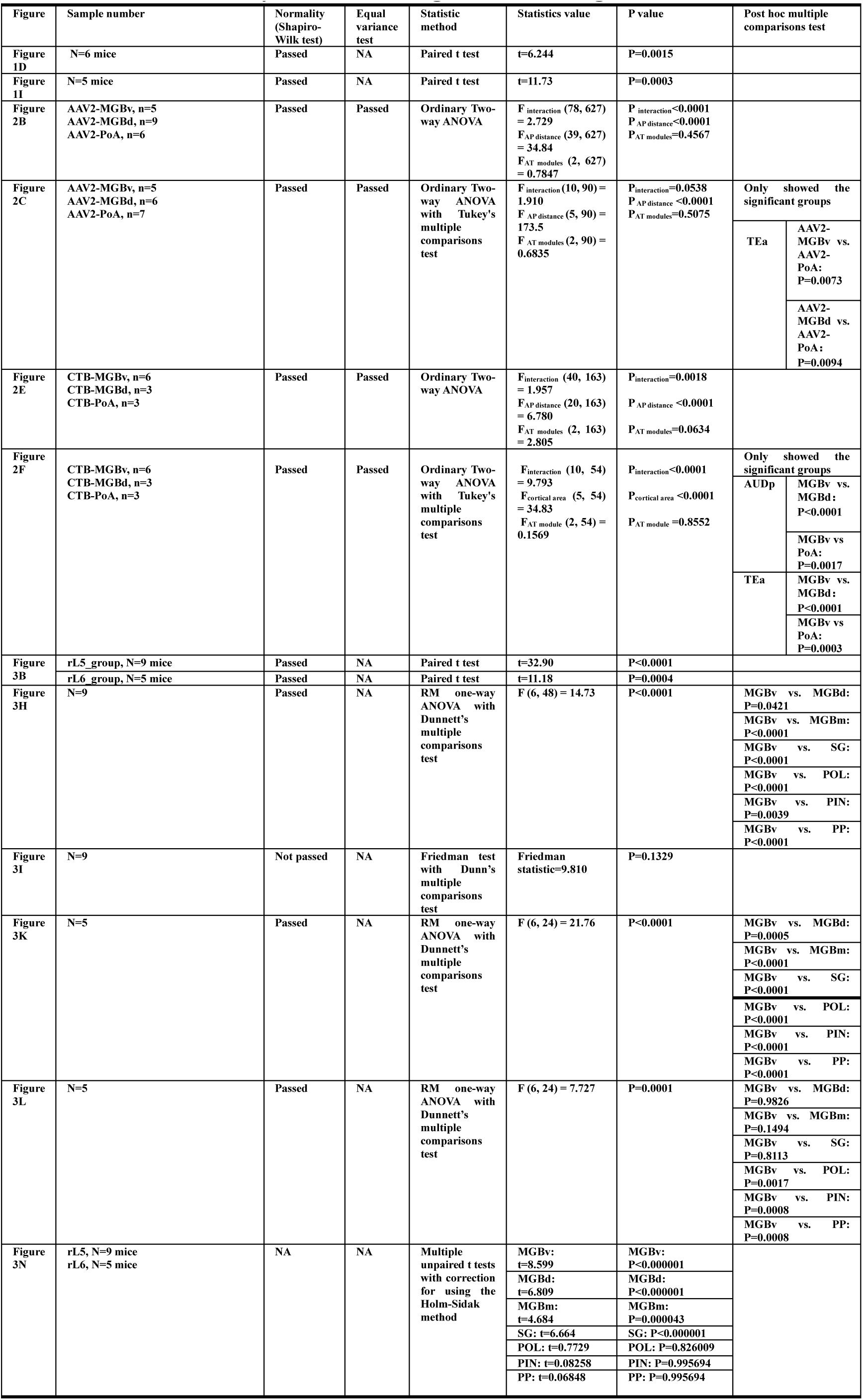
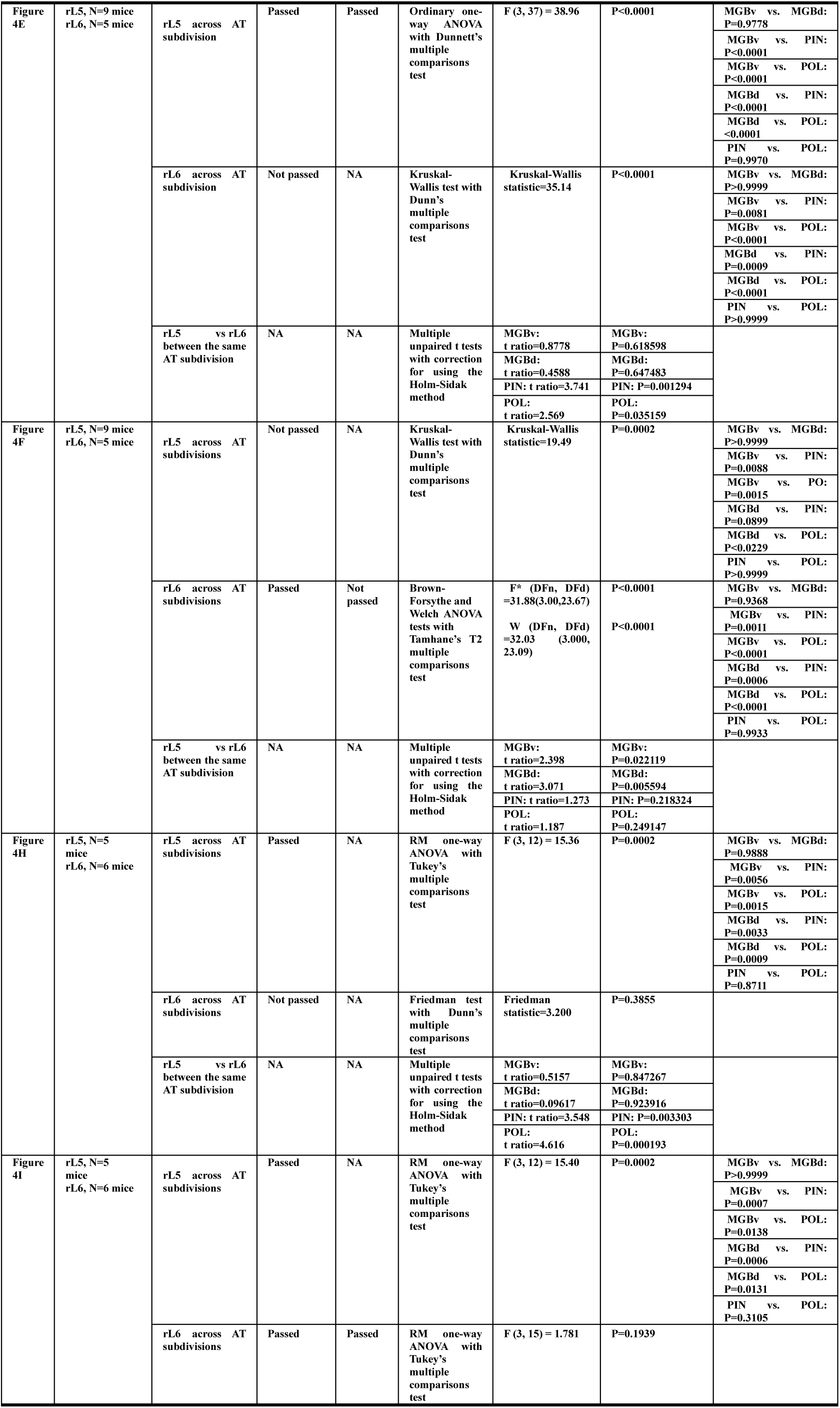
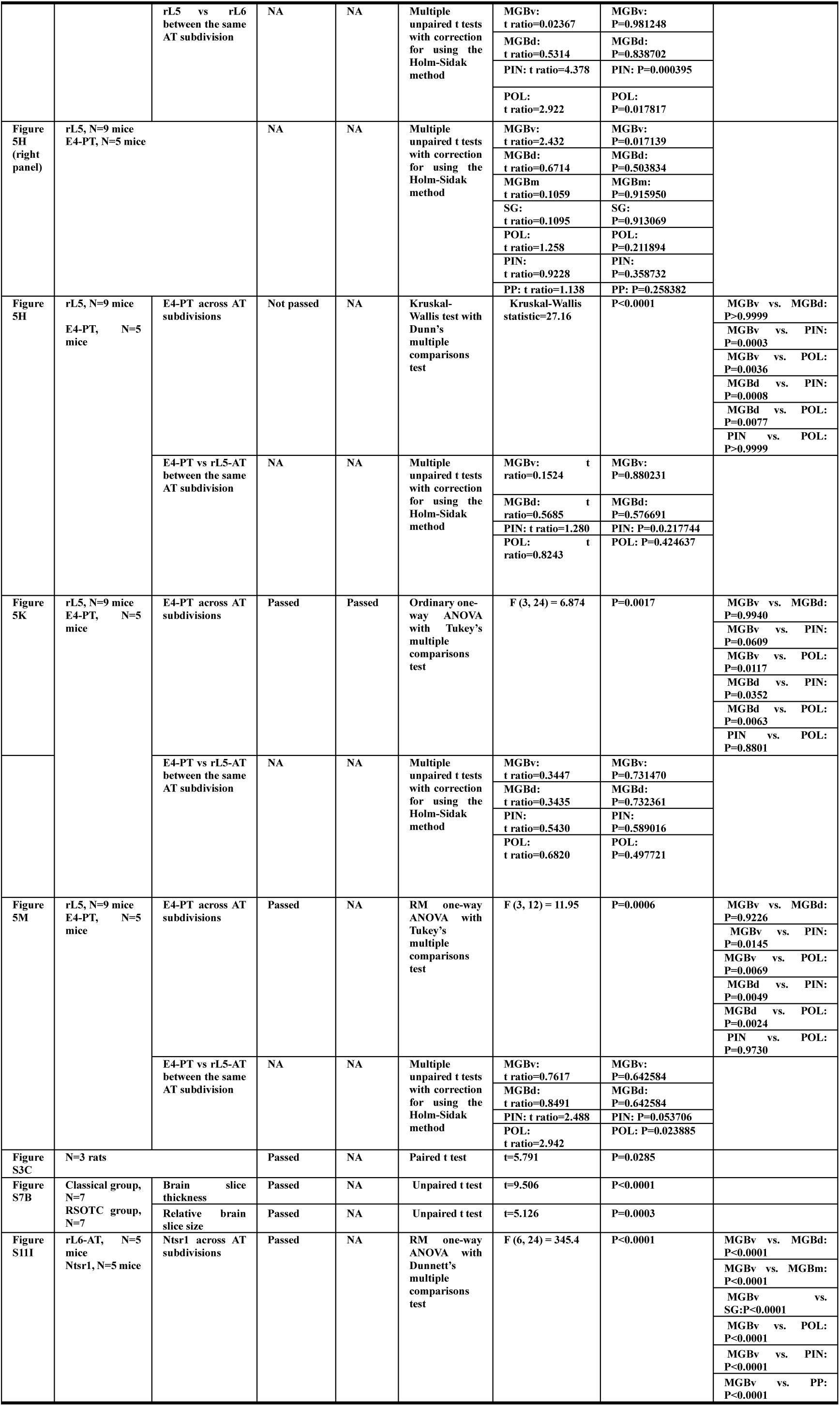
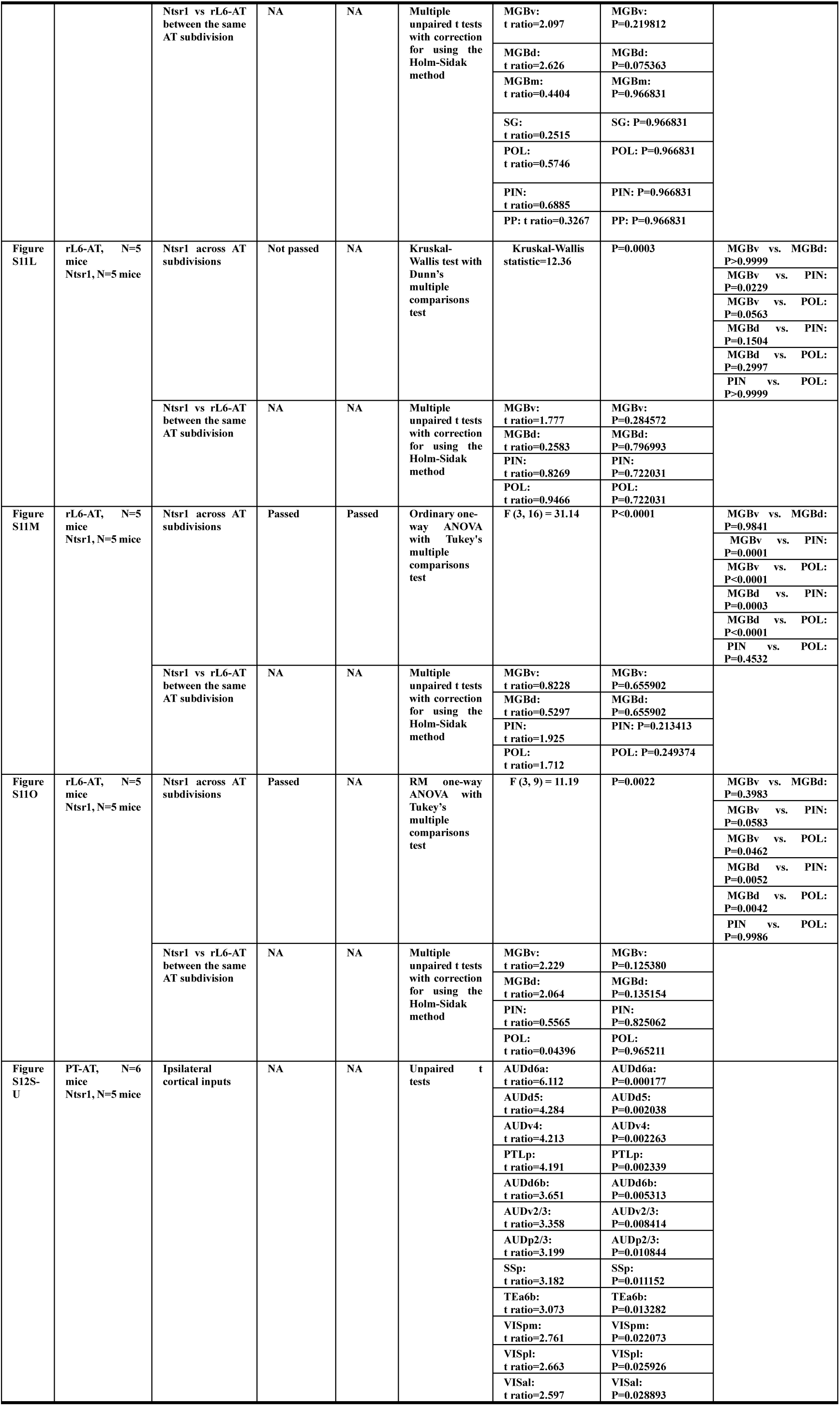
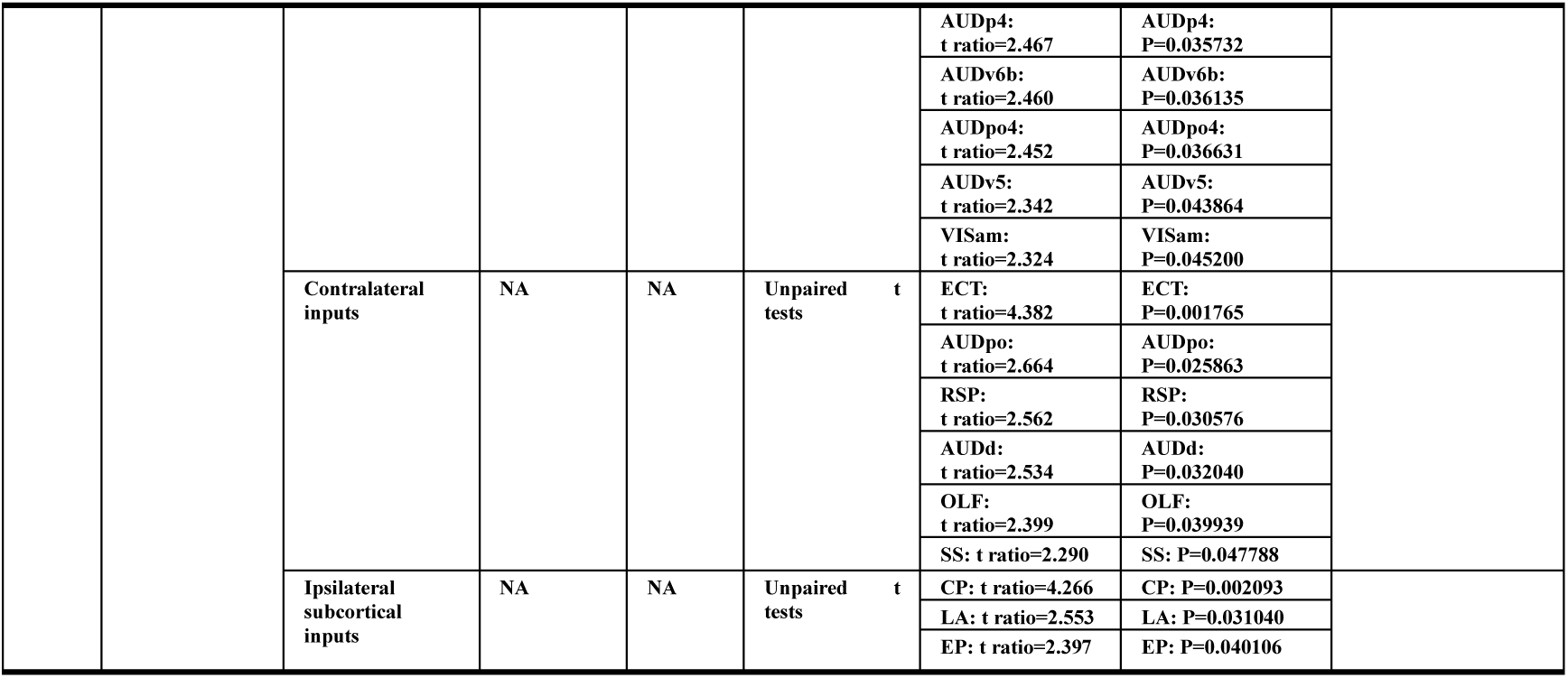
Statistical analysis relates to Figure 1-5 and Figure S1-S12.

